# ADGRG6 promotes adipogenesis and is involved in sex-specific fat distribution

**DOI:** 10.1101/2022.06.24.497411

**Authors:** Hai P. Nguyen, Aki Ushiki, Rory Sheng, Cassidy Biellak, Kelly An, Hélène Choquet, Thomas J. Hoffman, Ryan S. Gray, Nadav Ahituv

## Abstract

Fat distribution differences between males and females are a major risk factor for metabolic disease, but their genetic etiology remains largely unknown. Here, we establish *ADGRG6* as a major factor in adipogenesis and gender fat distribution. Deletion of *ADGRG6* in human adipocytes impairs adipogenesis due to reduced cAMP signaling. Conditionally knocking out *Adgrg6* in mouse adipocytes or deleting an intronic enhancer associated with gender fat distribution generates males with female-like fat deposition, which are protected against high-fat-diet-induced obesity and have improved insulin response. To showcase its therapeutic potential, we demonstrate that CRISPRi targeting of the *Adgrg6* promoter or enhancer prevents high-fat-diet-induced obesity. Combined, our results associate *ADGRG6* as a gender fat distribution gene and highlight its potential as a therapeutic target for metabolic disease.

## Introduction

Obesity, an excess of white adipose tissue (WAT), is a global epidemic and is closely associated with chronic metabolic diseases, such as type 2 diabetes and cardiovascular disease (*1, 2*). Obesity-related metabolic diseases are not simply the result of excess fat, but rather the distribution of adipose tissue has a major effect on these co-morbidities. While most studies have used clinical measures of overall fat and body mass index (BMI) to estimate disease risk, many studies using CT and MRI clearly show that abdominal visceral adipose tissue (VAT), but not subcutaneous adipose tissue (SAT), is associated with an increased risk for type 2 diabetes (*3-9*). Additional studies also showed that increased VAT represents a risk factor for developing insulin resistance and cardiovascular disease (*4-6, 10-15*). During periods of high metabolic activity, VAT release free fatty acids (FFA) that contribute more to plasma FFA than SAT (*12*). As a less metabolic active organ, SAT has better long-term lipid storage capacity and is a buffer during intake of dietary lipids, protecting other tissues from lipotoxic effects (*16*). In addition, SAT was found to be associated with increased HDL and decreased LDL cholesterol levels (*17-19*), indicating its protective role for cardiovascular disease. Upon cold or b-adrenergic stimuli, SAT can also acquire characteristics of brown adipose tissue (BAT), known as browning (*20-23*), becoming more metabolically active by dissipating energy via thermogenesis. Thus, increased SAT with browning potential correlates with improved insulin sensitivity (*24-26*).

Body fat distribution significantly differs between genders. Fat distribution differences are observed before puberty but become more prominent upon puberty (*27, 28*). Following puberty, females predominantly accumulate SAT, while males amass significantly more VAT. Women accumulate fat in the hip and limbs, while men accumulate a greater extent of fat in the trunk (*29-31*). Accumulation of adipose tissue around the viscera, the body’s internal organs, is associated with an increased risk of disease in both men and women (*3-5, 32*). In contrast, the preferential accumulation of adipose tissue in the lower extremities, such as the hips and legs, has been suggested to contribute to a lower incidence of myocardial infarction and coronary death in women during middle age (*33, 34*). This difference is thought to be due to numerous genes that are differentially expressed in adipose tissue between obese males and females, with only a few located on sex chromosomes (*35*). Many of these genes are involved in immune response and lipid and carbohydrate metabolism as well as clock genes, including *PER2, BMAL2*, and *CRY1* (*36*). The differential distribution of body fat between genders has been attributed to downstream effects of sex hormone secretion. Although sex steroids, especially estrogen, are involved in determining adipose distribution, several additional factors also play an important role. Recent human genome-wide association studies (GWAS) have identified multiple novel loci and pathways associated with measures of central obesity (*37, 38*). A recent GWAS study on body fat distribution difference between genders, identified multiple loci that are gender-heterogenous and associated with gender-specific fat distribution (*39*). Many of these loci have marked gender dimorphic patterns, the majority of which have stronger associations in women than in men; however, the mechanisms underlying this dimorphism remain largely unknown.

In a GWAS for gender-specific fat distribution in the UK Biobank, a noncoding single nucleotide polymorphism (SNP), rs6570507, near the adhesion G protein-coupled receptor G6 (*ADGRG6*; also called *GPR126*) showed gender heterogeneity for trunk fat ratio association, and was found to be associated with female trunk fat, but not with males (*39*). *ADGRG6* is a G-coupled receptor that is involved in the formation of the myelin sheath, regulates Schwann cell differentiation via activation of cyclin adenosine monophosphate (cAMP) (*40-42*), and maintains connective tissue in intervertebral disc (*43, 44*), inner ear (*45*), ventricles (*46*), and placenta (*47*). Ablation of *Adgrg6* in mouse 3T3-L1 adipocytes has been shown to prevent adipocyte differentiation (*48*). The *ADGRG6* locus is also associated with adolescent idiopathic scoliosis (AIS), and several enhancers in this locus were previously characterized by our lab due to this association (*49*). However, there is very little known about the role of *ADGRG6* in adipose tissue. Here, we performed gender-specific genetic association analyses using the UK Biobank (*50, 51*) and found that rs9403383, which is in high linkage disequilibrium (LD) with rs6570507 (r^2^=0.99), is associated with female trunk fat (P-value= 5.03478e-13) but is not associated with trunk fat in men. We show that rs9403383, leads to reduced enhancer activity and via transcription factor binding site and chromatin immunoprecipitation analyses demonstrate that the associated SNP affects HoxA3, GR (glucocorticoid receptor), and PGR (progesterone receptor) binding to an enhancer region. Knockout of this gene or enhancer in human adipocytes leads to impaired adipogenesis. Conditional knockout of *Adgrg6* in adipocytes in mice, using two different *Cre* driving promoters (*Pdgfra* and *Fabp4*), leads to fat deposition differences, making males more female like, protects against high-fat-diet-induced obesity, and shows improved glucose tolerance and insulin sensitivity. Furthermore, removal of the adipocyte enhancer in mice similarly leads to female-like fat deposition, lower body weight in male mice, high-fat-diet induced obesity protection and improved insulin response. Finally, CRISPRi targeting of the promoter or enhancer of *Adgrg6* prevents high-fat-diet induced obesity and improves insulin response. Combined, our results identify *ADGRG6* as an important adipogenesis factor regulating gender fat deposition and showcase its use as a therapeutic target to treat obesity and its co-associated morbidities.

## Results

### A preadipocyte enhancer near *ADGRG6* is associated with gender-specific fat distribution

A SNP in an intron of *ADGRG6*, rs6570507, was found to be associated with gender body fat distribution from segmental bioelectrical impedance analysis (sBIA) data (*39*), having a larger effect in female VAT than males. In addition, rs6570507 is also associated with body fat distribution using waist-to-hip ratio adjusted for BMI and obesity in females (*37, 52*). We examined *ADGRG6* expression in the Genotype-Tissue Expression (GTEX) portal (*53*), finding higher expression in males versus females in VAT (**fig. S1A**). rs9403383, which is in high LD with rs6570507 (r^2^=0.99), was found to be significantly associated with trunk fat mass (p-value=2.61×10^−11^) in the UK Biobank (*50, 51*) (**fig. S1B**) and to have significant *cis*-eQTL association with both VAT (p-value=3.26×10^−5^) and SAT in the GTEX portal (p=5.2×10^−13^) (**fig. S1C**). To determine whether SNP rs9403383 at *ADGRG6* was associated with trunk fat mass in humans and had different effects in men and women, we conducted sex-specific analyses in the European ancestry sample of the UK Biobank (UKB) cohort. While SNP rs9403383 was genome-wide significantly (beta [95% CI] = -0.016, [-0.021 to -0.012]; *P*-value= 5.03 × 10^−13^) with trunk fat mass in women, we observed no significant association (beta [95% CI] = -0.0044 [-0.0044 to 0.0002]; *P*-value = 0.06) in men (N=205,628). In addition, analysis of the Genome Aggregation Database (gnomAD) (version 3.1.2) of control/biobank populations (*54*), we found that rs9403383 has higher allele frequency in females (0.4618) than in males (0.3951).

We next set out to assess whether rs6570507 or SNPs in LD with it (r^2^>0.8) overlap potential active enhancer sequences, by analyzing H3K27ac ChIP-seq data from human adipose tissue (*55*). We found two H3H27ac peaks, which were previously named in a previous study in our lab that characterized AIS-associated enhancers, named *ADGRG6_3* and *ADGRG_4* (*49*), to overlap rs6570507 and rs9403383, respectively (**Fig.1A**). To functionally validate whether sequences at both H3K27ac peaks might have adipocyte enhancer activity, we cloned both regions into an enhancer assay vector, pGL4.23 (Promega), that contains a minimal promoter followed by the luciferase reporter gene. We transfected these constructs into human preadipocytes, finding only *ADGRG6_4* to have enhancer activity, showing ∼15-fold higher luciferase expression than the negative control (**Fig. 1B**). We next tested whether rs9403383 affects *ADGRG6_4* enhancer activity, finding that the gender body fat distribution associated variant significantly reduced enhancer activity compared to the unassociated variant (**Fig. 1C**).

**Fig. 1.**
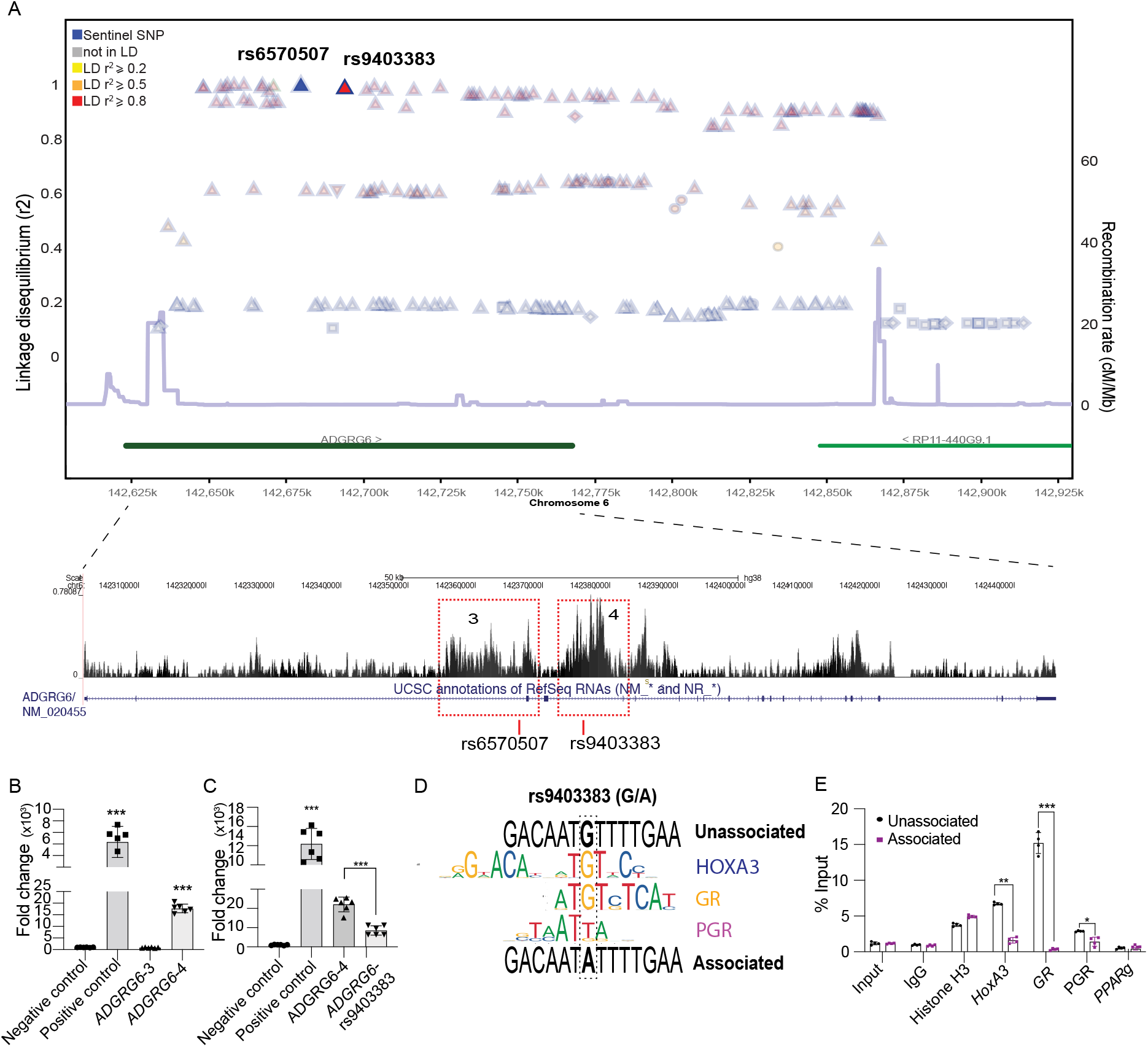
A preadipocyte enhancer in the intron of *ADGRG6* is associated with gender-specific fat distribution. (**A**) A linkage disequilibrium plot and integrative genomic viewer snapshot for the *ADGRG6* locus showing the lead SNP rs6570507 and the linked SNP rs9403383 with H3K27ac ChIP-seq peaks, *ADGRG6*_3 and *ADGRG6*_4, from human adipose tissue [49]. The colors of the triangles reflect the degree of linkage disequilibrium (LD) with the lead SNP measured by r^2^ and the estimated recombination rates (cM/Mb) from HapMap CEU release 22 shown as gray lines. (**B**) Luciferase assays in human preadipocytes for *ADGRG6*_3 and *ADGRG6*_4. pGL4.13 (Promega) with an SV40 early enhancer was used as a positive control and the pGL4.23 empty vector as a negative control. Data are represented as mean ± S.D ***≤0.001. (**C**) Luciferase assays for *ADGRG6*_4 and the associated variant containing rs9403383. Data are represented as mean ± S.D ***≤0.001. (**D**)Transcription factor binding site analysis of unassociated and associated variant sequences showing HOXA3, GR, and PGR binding sites. (**E**) HOXA3, GR, and PGR ChIP-qPCR relative to input for the unassociated and associated variant. IgG and PPARg are used as negative controls and Histone H3 as a positive control. Data are represented as mean ± S.D *≤0.05, **≤0.01, ***≤0.00

We next examined whether rs9403383 leads to transcription factor binding site (TFBS) changes. Using PROMO (*56*) and TRANSFAC (*57, 58*), we found that the unassociated sequence contains TFBS for multiple TFs that are critical for adipocyte differentiation, such as CCAAT/enhancer-binding protein delta (C/EBPd) and glucocorticoid receptor (GR). The associated variant could disrupt the TFBS of Homeobox protein A3 (HoxA3), progesterone receptor (PGR) and GR (**Fig. 1D**). We generated a human preadipocyte cell line carrying the rs9403383 associated variant by transfecting Cas9, gRNA, and a ssDNA donor sequence containing the associated variant. We selected for single-cell colonies and validated them for the existence of this variant by sequencing. We then preformed ChIP-qPCR with either HOXA3, GR, or PGR antibodies on human preadipocytes containing either the unassociated or associated rs9403383 variant. We also used IgG and PPARg as negative controls and histone H3 as a positive control. We observed significantly decreased enrichment for all three transcription factors (HOXA3, GR, or PGR) in the cells having the associated allele (**Fig. 1E**), indicating that the associated rs9403383 variant affects the binding of these transcription factors to the enhancer sequence. Taken together, we identified a novel adipocyte enhancer in the intron of *ADGRG6* that carries a SNP associated with gender-specific fat distribution and can hinder its activity.

### *ADGRG6* is highly expressed in adipose progenitors and mesenchymal stem cells

We next examined the role of *ADGRG6* in adipose tissue. We measured, via qRT-qPCR, *ADGRG6* mRNA levels during human adipocyte differentiation and found them to be the highest in preadipocytes and to significantly decrease when subjected to adipocyte differentiation (**fig. S2A**). The expression pattern of *ADGRG6* was similar to *SOX9*, which is required for adipogenesis and is known to decrease during adipocyte differentiation (*59*). In contrast, *PPARg* and *FABP4*, known adipogenic markers, were significantly increased during adipogenesis (**fig. S2A**). Analysis of ADGRG6 protein levels using Western showed similar results, whereby its protein expression was higher in preadipocytes and markedly decreased upon adipocyte differentiation (**fig. S2A**).

We next examined *Adgrg6* expression levels in mouse adipose tissues. We isolated the stromal vascular fraction (SVF) and adipocytes of pWAT of male mice. Gene analysis revealed that *Adgrg6* mRNA levels were similar to *Sox9*, being 100-fold higher in the SVF fraction than adipocytes and in contrast to *Fabp4* levels which were significantly higher in the adipocyte fraction (**fig. S2B**). We next FACS-isolated mesenchymal stem cells (MSC) (CD105+), adipose progenitors (APC) (CD34+, Pdfrga+) and immune cells (CD45+) from the SVF. We observed that *Adgrg6* is highly expressed in MSC, indicating that Adgrg6 might be required for adipose lineage development (**fig. S2C**). Overall, these results demonstrate that *ADGRG6* is highly expressed in early adipose progenitors and MSCs and significantly decreases in expression during adipocyte differentiation, potentially playing an early role in adipogenesis. In addition, *Adgrg6* shows a similar expression pattern in both humans and mice (**fig.S2D**), suggesting that mice could be used as a model to characterize its function.

### *ADGRG6* is involved in adipogenesis

To characterize the role of *ADGRG6* in adipogenesis, we knocked out the gene, the *ADGRG6_4* enhancer, and generated a human preadipocyte line containing the associated rs9403383 SNP (**Fig. 2A**). For the gene knockout (KO), we transfected human preadipocytes with Cas9 protein and two gRNAs targeting exon 2 of *ADGRG6*. For the *ADGRG6_4* enhancer KO (EKO), human preadipocytes were transfected with gRNAs targeting the two ends of the enhancer sequence, along with Cas9 protein. For the associated SNP knockin (AKI), we first sequenced the human preadipocyte cell line and observed that it contains the unassociated allele. These human preadipocytes were transfected with a gRNA targeting *ADGRG6_4* and an ssDNA donor containing the associated SNP rs9403383. Single-cell clones for all three manipulations were FACS isolated and screened by genotyping for homozygous colonies. We then measured, using qRT-PCR, *ADGRG6* mRNA levels finding the gene KO cells to have nearly depleted levels of *ADGRG6* transcripts, EKO cells 50% reduction and the associated variant 20% reduction compared to wild-type (WT) cells (**Fig. 2B**). We then subjected the *ADGRG6* KO, *ADGRG6* EKO, associated variant and WT cells to adipocyte differentiation. mRNA expression analyses using qRT-PCR showed that *ADGRG6* KO, *ADGRG6_4* EKO, and *ADGRG6* AKI had lower expression of adipogenic markers, such as *C/EBPb, PPARg*, and *FABP4* compared to the WT cells. In contrast, *SOX9* levels, an adipogenesis inhibitor (*59*), were higher in *ADGRG6* KO, EKO, AKI compared to WT cells (**Fig. 2B**). Oil red O lipid staining showed that KO, EKO, and AKI cells have lower lipid accumulation compared to WT (**Fig. 2C**). In addition, using Fiji (*60*), to quantify lipid droplet size, we observed *ADGRG6* KO, *ADGRG6_4* EKO, and *ADGRG6* AKI cells to have more smaller lipid droplets containing cells than WT cells (**Fig. 2C**).

**Fig. 2.**
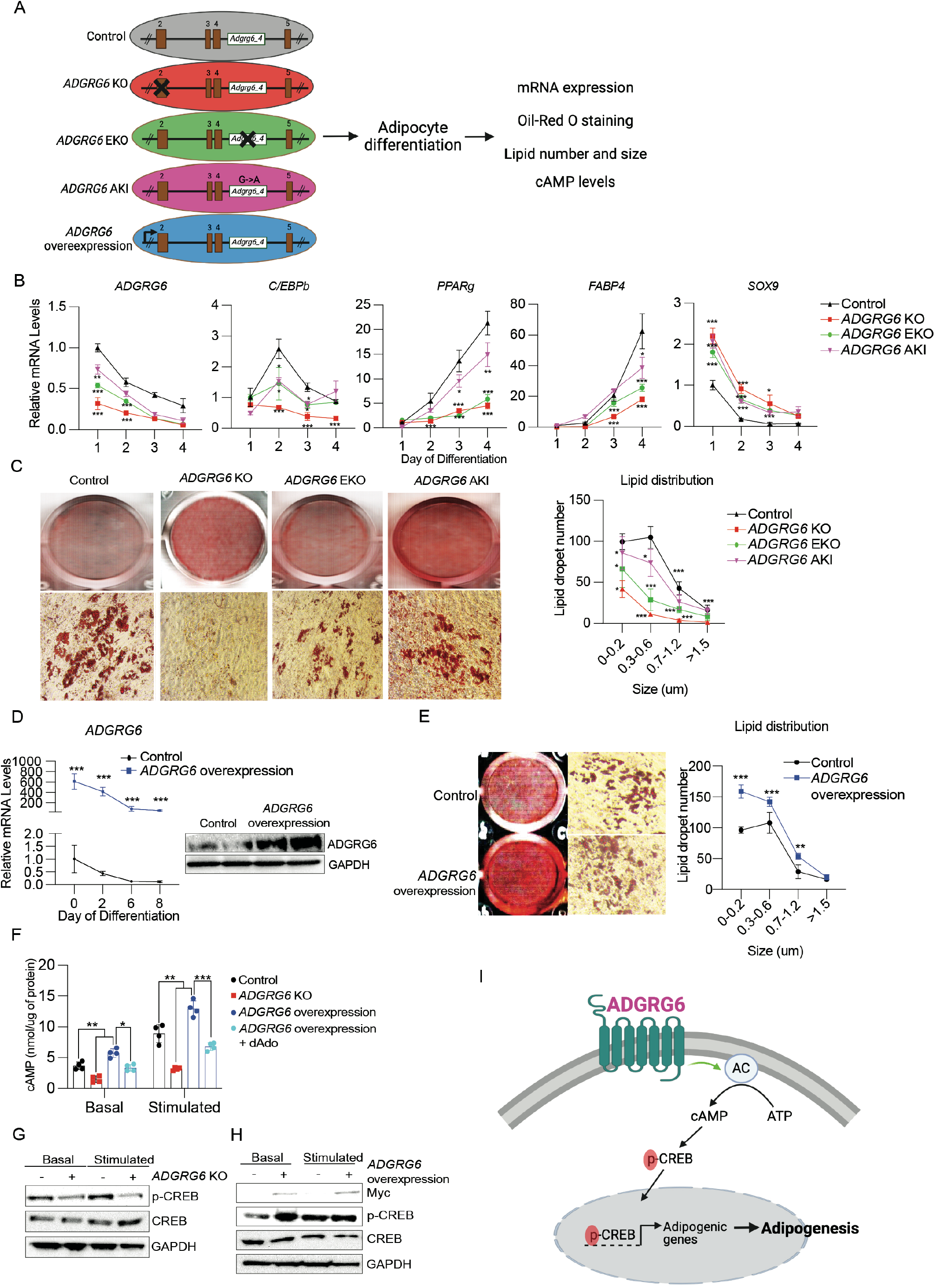
*ADGRG6* regulates adipogenesis via cAMP. (**A**) Schematic of the various *ADGRG6* manipulations generated in human preadipocytes. *ADGRG6* knockout (KO), enhancer knockout (EKO), *ADGRG6* associated knockin (AKI) and *ADGRG6* overexpression. (**B**) qRT-PCR of *ADGRG6* and adipogenic markers (*C/EBPb, PPARg, FABP4* and *SOX9*) during adipocyte differentiation of control, *ADGRG6* KO, *ADGRG6* EKO, and *ADGRG6* AKI cells. Data are represented as mean ± S.D *≤0.05, **≤0.01, ***≤0.001. (**C**) Oil red O staining (left) and lipid distribution (Right) by Fiji of control, *ADGRG6* KO, *ADGRG6* EKO, and *ADGRG6* AKI cells. Data are represented as mean ± S.D *≤0.05, ***≤0.001. (**D**) qRT-PCR of *ADGRG6* during adipocyte differentiation of control and *ADGRG6* overexpressing cells (Left). Immunoblotting for ADGRG6 (Right). Data are represented as mean ± S.D ***≤0.001. (**E**) Oil red O staining (left) and lipid distribution (Right) by Fiji of control, *ADGRG6* overexpressing cells. Data are represented as mean ± S.D **≤0.01, ***≤0.001. (**F**) cAMP levels of control, *ADGRG6* KO, *ADGRG6* overexpressing cells and *ADGRG6* overexpressing cells treated with 2’,5’-Dideoxyadenosine (dAdo) in basal condition or stimulated with forskolin. Data are represented as mean ± S.D *≤0.05, **≤0.01, ***≤0.001. (**G**) Immunoblotting for phospho-CREB, CREB, and GAPDH in *ADGRG6* KO in basal condition or stimulated with forskolin. (**H**) Immunoblotting for myc-ADGRG6, phospho-CREB, CREB, and GAPDH in *ADGRG6* overexpressing cells in basal condition or stimulated with forskolin. (**I**) Proposed model of *ADGRG6* function during adipogenesis.

To analyze the effect of increased *ADGRG6* expression on adipogenesis, we overexpressed *ADGRG6* in human preadipocytes. Myc-tagged *ADGRG6* was transfected into human preadipocytes, followed by their differentiation into adipocytes. *ADGRG6* mRNA levels, as measured by qRT-PCR, were significantly higher in the overexpressing cells compared to WT cells throughout differentiation (**Fig. 2D**). Immunoblotting further confirmed the overexpression of *ADGRG6* (**Fig. 2D**). In contrast to the knockout cells, *ADGRG6* overexpressing cells exhibited higher levels of adipogenic genes (*C/EBPb, C/EBPd* and *FABP4*) and lower levels of *SOX9* compared to WT cells (**fig. S3**). In addition, *ADGRG6* overexpressing preadipocytes had much higher lipid accumulation than control cells and increased lipid droplet size (**Fig. 2E**). Combined, these data suggest that *ADGRG6* has an important role in adipogenesis and deletion of its enhancer or the fat distribution associated SNP reduces its expression level leading to impaired adipogenesis.

### *ADGRG6* increases cAMP levels during adipogenesis

Previously characterized ADGRG6 ligands, such as Type IV collagen and Laminin-211, were shown to induce an adenosine 3’-5’-monophosphate (cAMP) response (*42, 61*). cAMP is a well-known signaling molecule that initiates adipogenesis by increasing protein kinase A (pKA), which then phosphorylates early transcription factors, including *C/EBPb* (*62-64*). To test whether ADGRG6 might increase cAMP levels in preadipocytes, we measured cAMP levels in WT and *ADGRG6* KO human preadipocytes in both basal condition and treatment with forskolin, which is an agonist pKA, using ELISA. We found that cAMP levels were approximately 2-fold lower in *ADGRG6* KO cells compared to WT cells. In addition, upon forskolin treatment, WT cells had a 2-fold increase in cAMP levels compared to the basal condition while *ADGRG6* KO cells did not show a significant increase (**Fig. 2F**). We also measured cAMP levels in human preadipocytes that overexpress *ADGRG6*. In the basal condition, *ADGRG6* overexpression significantly increased cAMP levels and in the stimulated condition, cAMP levels were increased by 2-fold compared to WT (**Fig. 2F**). In order to test whether *ADGRG6* can directly affect cAMP levels via adenyl cyclase (AC), that converts ATP to cAMP, we treated *ADGRG6* overexpressing preadipocytes with 2’,5’-Dideoxyadenosine (aAdo), an inhibitor of AC. In the basal condition, cAMP levels were reduced in both WT and *ADGRG6* overexpressing cells upon aAdo treatment (**Fig. 2F**). Moreover, in the Ado stimulated condition, the *ADGRG6* overexpressing preadipocytes only increased cAMP levels by 25% compared to WT (**Fig. 2F**). Furthermore, we also examined the downstream target of cAMP-PKA, CREB phosphorylation. Immunoblotting revealed that in basal condition, *ADGRG6* KO cells have lower levels of p-CREB compared to control cells (**Fig. 2G**), while *ADGRG6* overexpression significantly increased p-CREB (**Fig. 2H**). In the stimulated condition, p-CREB increased in WT cells but not in *ADGRG6* KO cells (**Fig. 2G**). In contrast, *ADGRG6* overexpression did not further increase p-CREB (**Fig. 2H**), likely due to significant increase in the basal condition. Taken together, our result suggests that ADGRG6 increases cAMP levels and its downstream target CREB in human preadipocytes to promote adipocyte differentiation (**Fig. 2I**).

### Conditional knockout of *Adgrg6* in adipocytes leads to female-like fat depots in males

To examine the role of *Adgrg6* on adipose tissue development *in vivo*, we generated adipose-specific knockout mice (*Adgrg6*^*ASKO*^). We crossed *loxP* flanked *Adgrg6* exon 3 and 4 (*Adgrg6*^*fl/fl*^) mice (*65*) with the platelet derived growth factor receptor alpha (*Pdgfra*) promoter-driven *Cre* mice (*Pdgfra-Cre*), in which Cre is highly expressed in adipose-lineage cells and progenitors (*66*) (**Fig. 3A**). qRT-PCR analysis of *Adgrg6*^*ASKO/ASKO*^ homozygous mice, showed *Adgrg6* mRNA levels to be 80% in iWAT (inguinal) and 40% in pWAT (perigonadal) compared to the Cre(-)/floxed *Adgrg6* mice (termed hereafter as control mice). This ablation efficiency is reported in previous studies using the *Pdgfra-Cre* mice (*67*). Notably, there was no significant difference in *Adgrg6* expression between female *Adgrg6*^*ASKO/ASKO*^ and female control littermates (**Fig. 3B**). Adipogenic markers, such as *Pparg* and *Fabp4* were decreased in iWAT and pWAT in *Adgrg6*^*ASKO/ASKO*^ male mice and *Sox9*, an adipogenesis inhibitor, showed higher mRNA levels compared to control littermate male mice (**fig. S4A**). *Adgrg6*^*ASKO/ASKO*^ male mice showed significantly lower body weight (BW) than WT mice, while the female *Adgrg6*^*ASKO/ASKO*^ mice displayed no difference in BW compared to their control littermates (**Fig. 3C**). There was no difference in food intake between control and KO mice (**fig. S4B**). However, the BW of the *Adgrg6*^*ASKO/ASKO*^ male mice was similar to age-matched female KO mice (**Fig. 3C**).

**Fig. 3.**
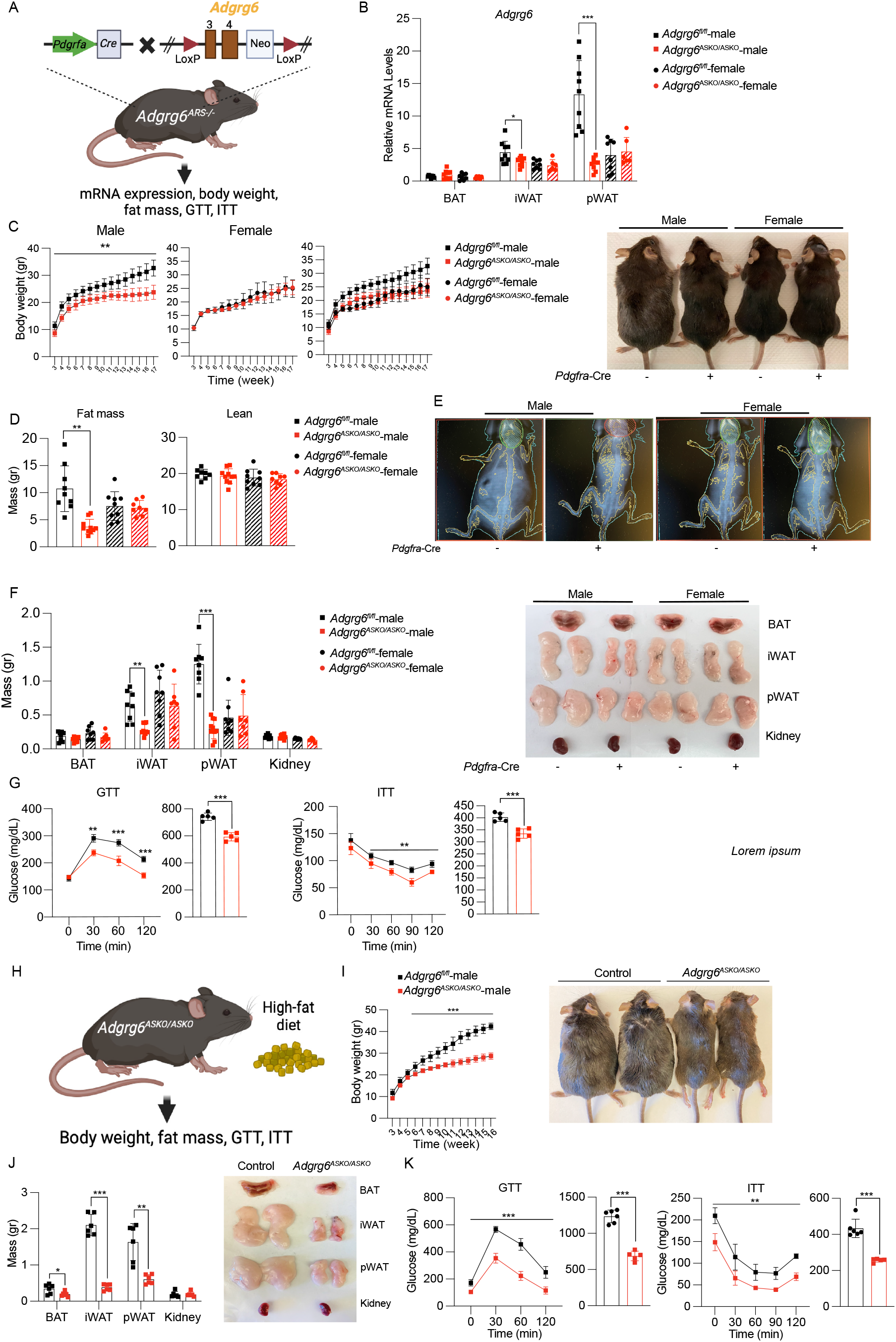
*Adgrg6* adipose-specific knockout leads to female-like fat distribution in male mice. (**A**) Schematic of *Adgrg6* adipose-specific knockout mice (*Adgrg6*^*ASKO/ASKO*^) generation and a summary of the measured metabolic phenotypes. Glucose tolerance test (GTT), insulin tolerance test (ITT). (**B**) qRT-PCR of *Adgrg6* in brown adipose tissue (BAT), inguinal adipose tissue (iWAT), and perigonadal adipose tissue (pWAT) of control *Adgrg6*^*fl/fl*^ (male=9 mice, female=8) and *Adgrg6*^*ASKO/ASKO*^ (male=9, female=6) mice. Data are represented as mean ± S.D *≤0.05, **≤0.01, ***≤0.001. (**C**) Body weight (left) of control (male=9 mice, female=8) and *Adgrg6*^*ASKO/ASKO*^ (male=9, female=8) mice measured for 17 weeks. Data are represented as mean ± S.D **≤0.01. Representative mouse image of mice at 17-week-old and fed on chow diet (right) (**D**) Fat and lean mass of control (male=9 mice, female=9) and *Adgrg6*^*ASKO/ASKO*^ (male=10, female=8) measured by dual energy X-ray absorptiometry (DEXA). Data are represented as mean ± S.D **≤0.01. (**E**) Whole-body scan images of control and *Adgrg6*^*ASKO/ASKO*^ male and female mice. (**F**) Tissue weight (left) of control (male=8 mice, female=8) and *Adgrg6*^*ASKO/ASKO*^ (male=10, female=7). Data are represented as mean ± S.D **≤0.01, ***≤0.001. Representative tissue images (right) of mice fed on regular chow diet at 17-weeks of age. (**G**) Glucose tolerance test (GTT) and insulin tolerance test (ITT) of male control (n=5 mice) and *Adgrg6*^*ASKO/ASKO*^ (n=5). Data are represented as mean ± S.D **≤0.01, ***≤0.001. (**H**) Schematic of male *Adgrg6*^*ASKO/ASKO*^ mice fed with high-fat diet (HFD) and the metabolic phenotypes measured. (**I**) Body weight (left) of control and *Adgrg6*^*ASKO/ASKO*^ male mice (N=6) on HFD measured for 16 weeks. Data are represented as mean ± S.D ***≤0.001. Representative mouse images (right) of 20-week-old mice that were fed on HFD for 16 weeks. (**J**) Tissue weight (left) of male control (n=6 mice) and *Adgrg6*^*ASKO/ASKO*^ (n=5) mice. Data are represented as mean ± S.D **≤0.01, ***≤0.001. Representative tissue images (right) from 20-week-old mice fed on HFD for 16 weeks. (**K**) Glucose tolerance test (GTT) with area under the curve (AUC) and insulin tolerance test (ITT) with AUC of male control (n=6) and *Adgrg6*^*ASKO/ASKO*^ (n=5) 20-week-old mice fed on HFD for 16 weeks. Data are represented as mean ± S.D *≤0.05 **≤0.01, ***≤0.001.

The *Pdgfra*-driven promoter has been utilized to label and genetically modify adipose lineage genes; however, its expression is also detected in other tissues, such as the retina and glial cells (*66-69*). We thus also conditionally knocked out *Adgrg6* by crossing *Adgrg6*^*fl/fl*^ mice with the adiponectin C1Q and collagen domain containing (*Adipoq*) promoter-driven Cre mice (*Adipoq*-*Cre*). The *Adipoq* promoter is known to drive expression in more mature adipocytes (*70*); however some studies utilized this mouse model to manipulate genes expressed in earlier adipose progenitors (*71, 72*). Similar to the *Pdgfra*-Cre driven mice, homozygous *Adipoq-Cre;Adgrg6*^*fl/fl*^ male mice showed reduced body weight compared to control mice (**fig. S4D**). While the effect on BW was not as substantial as that of the *Pdgfra*-driven Cre, likely due to *Adgrg6* expression being primarily in early adipose progenitors, the difference remained significant. All subsequent experiments were carried out in *Pdgfra*-driven Cre mice due to their stronger adipose progenitor expression.

We next used dual energy X-ray absorptiometry (DEXA) to examine body fat mass in these mice. We observed that the fat mass of *Adgrg6*^*ASKO/ASKO*^ male mice was approximately six grams lower than control littermates, with no difference in lean mass. There was no difference in fat mass between *Adgrg6*^*ASKO/ASKO*^ female mice and the perspective female WT mice (**Fig. 3D**). *Adgrg6*^*ASKO/ASKO*^ male mice displayed similar fat mass to age-matched control female mice (**Fig. 3D**). While *Pdgfra* is not thought to be involved in skeletal development (*73*) and *Adgrg6* does play a role in this process (*65*), but was associated with AIS, we assessed if skeletal development was altered. Analysis of spine morphology and bone mineral density (BMD) did not find any apparent skeletal abnormalities (**Fig. 3E**) nor altered BMD (**fig. S4C**) in *Adgrd6*^*ASKO/ASKO*^ mice. Upon tissue dissection, *Adgrg6*^*ASKO/ASKO*^ male mice showed significantly lower iWAT and pWAT mass compared to control littermates, with no changes in brown adipose tissue (BAT) or kidney (**Fig. 3F**). No significant differences were observed between female *Adgrg6*^*ASKO/ASKO*^ and control littermates (**Fig. 3F**). As the BW of *Adgrg6*^*ASKO/ASKO*^ male mice was reduced and similar to control females, we next examined their glucose tolerance (GTT) and insulin sensitivity (ITT). We observed that *Adgrg6*^*ASKO/ASKO*^ male mice have significantly higher glucose tolerance than control mice at all measured time points, with significantly improved insulin sensitivity (**Fig. 3G**).

To further evaluate *Adgrg6* ablation on adiposity, we placed *Adgrg6*^*ASKO/ASKO*^ male mice on a high-fat diet (HFD) (**Fig. 3H**). Compared to control mice, *Adgrg6*^*ASKO/ASKO*^ male mice gained significantly less BW (**Fig. 3I**). In addition, the weight of all adipose depots, including BAT, iWAT, and pWAT of *Adgrg6*^*ASKO/ASKO*^ male mice was significantly lower than control mice with no changes in the kidney (**Fig. 3J**). In addition, *Adgrg6*^*ASKO/ASKO*^ male mice showed significantly improved GTT and ITT compared to control males (**Fig. 3K**), indicating that *Adgrg6* ablation in males can reduce diet-induced obesity and improve obesity-associated insulin resistance. Taken together, our data suggest that *Adgrg6* has an important role in adipogenesis *in vivo* and its removal in adipocytes leads to a female-like fat distribution in male mice, protecting them against HFD-induced obesity and providing improved insulin response.

### *Adgrg6_4* enhancer knockout male mice have female-like fat depots

We next set out to characterize the *in vivo* function of the *Adgrg6_4* enhancer by knocking it out in mice. We first identified sufficient mouse sequence homology to the human *Adgrg6_4* enhancer using LiftOver (*74*) and ENSEMBL (*75*) (**fig. S5A**). We designed two gRNA to target the enhancer and generated *Adgrg6_4* enhancer KO mice, which were named *Adgrg6*^*ARS-/-*^ (adipose regulatory sequence) using the improved-Genome editing via Oviductal Nucleic Acids Delivery (i-GONAD) approach (*76*) (**Fig. 4A**). Knockout mice were validated for proper targeting via genotyping and Southern blot (**fig. S5B**). Interestingly, analysis of *Adgrg6* mRNA expression levels by qRT-qPCR, found it to be significantly reduced only in pWAT of male *Adgrg6*^ARS-/-^ mice (**Fig. 4B**). There were no significant changes in *Adgrg6* expression in other adipose depots, including BAT and iWAT, and other tissues, such as liver and muscle. Moreover, *Adgrg6* mRNA levels in female *Adgrg6*^*ARS-/-*^ mice did not change (**Fig. 4B**). These results suggest that the enhancer of *Adgrg6* is only active in pWAT. Analysis of the mRNA expression of adipogenic markers, *C/ebpb, Pparg*, and *Fabp4* and the adipogenesis inhibitor, *Sox9*, showed changes similar to the conditional gene knockout only in pWAT of male *Adgrg6*^*ARS-/-*^ mice (**Fig. 4C**).

**Fig. 4.**
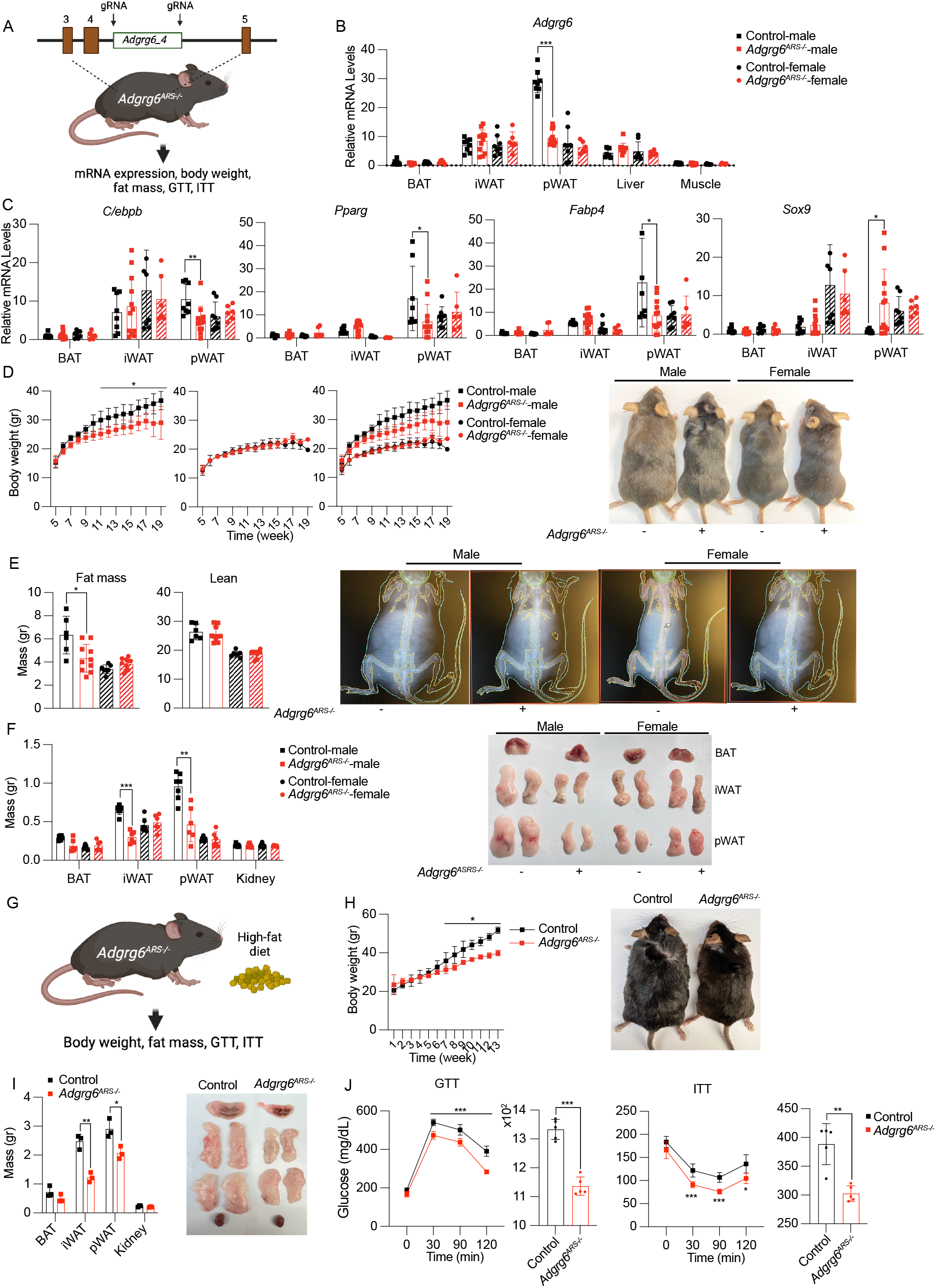
*Adgrg6* enhancer knockout exhibits gender-specific fat distribution. (**A**) Schematic of *Adgrg6_4* adipose enhancer knockout (*Adgrg6*^*ARS-/-*^) generation and a summary of the phenotypes measured. (**B**) qRT-PCR of *Adgrg6* in brown adipose tissue (BAT), inguinal adipose tissue (iWAT), and perigonadal adipose tissue (pWAT) of control (male=8 mice, female=8) and Adgrg6^ARS-/-^ (male=12, female=6). Data are represented as mean ± S.D. ***≤0.001. (**C**) qRT-PCR of adipogenic markers (*C/EBPb, PPARg, FABP4* and *SOX9*) in BAT, iWAT, and pWAT control (male=8 mice, female=8) and *Adgrg6*^*ARS-/-*^ (male=12, female=6) mice. Data are represented as mean ± S.D. *≤0.05, **≤0.01. (**D**) Body weight (left) of control (male=8 mice, female=8) and *Adgrg6*^*ARS-/-*^ (male=12, female=6) measured for 17 weeks. Data are represented as mean ± S.D. *≤0.05. Representative mouse images (right) of 19-week-old mice fed on regular chow diet. (**E**) Fat and lean mass of control (male=6, female=7) and *Adgrg6*^*ARS-/-*^ (male=10, female=10) measured by dual energy X-ray absorptiometry (DEXA) (Left). Data are represented as mean ± S.D. *≤0.05. Representative images of whole-body scans of control and *Adgrg6*^*ARS-/-*^ male and female mice (right). (**F**) Tissue weight (left) of control (male=7, female=7) and *Adgrg6*^*ARS-/-*^ (male=6, female=6) mice. Data are represented as mean ± S.D. **≤0.01, ***≤0.001. Representative tissue images (right) of 16-week-old mice fed on regular chow diet. (**G**) Schematic of male *Adgrg6*^*ARS-/-*^ mice fed with high-fat diet (HFD) and the metabolic phenotypes measured (**H**) Body weight (left) of control and *Adgrg6*^*ARS-/-*^ male mice (n=6) on HFD measured for 13 weeks. Data are represented as mean ± S.D *≤0.05. Representative mouse images (right) of 18-week-old mice fed on HFD for 13 weeks. (**I**) Tissue weight (left) of male control (n=3 mice) and *Adgrg6*^*ARS-/-*^ (N=3) mice. Data are represented as mean ± S.D *≤0.05, **≤0.01. Representative tissue images (right) of 8-week-old mice fed on HFD for 13 weeks. (**J**) Glucose tolerance test (GTT) with area under the curve (AUC) and insulin tolerance test (ITT) with AUC of male control (n=6) and *Adgrg6*^ARS-/-^ (n=6) of 18-week-old mice fed on HFD for 13 weeks. Data are represented as mean ± S.D *≤0.05 **≤0.01, ***≤0.001.

Similar to *Adgrg6*^*ASKO/ASKO*^ mice, male *Adgrg6*^*ARS-/-*^ mice exhibited lower body weight than littermate control mice, while there was no difference in female mice (**Fig. 4D**). Using DEXA, we showed that the fat mass of male *Adgrg6*^*ARS-/-*^ mice was approximately two grams lower than control male mice and similar to female mice (**Fig. 4E**). There was no difference in lean mass. Evaluation of bone mass and density due to *Adggr6* known role in chondrocyte development found no changes in skeletal abnormalities in *Adgrg6*^*ARS-/-*^ mice (**Fig. S5C**). We next measured the mass of all adipose depots. We observed that male *Adgrg6*^*ARS-/-*^ mice had lower pWAT and iWAT mass (**Fig. 4F**). The lower iWAT mass could be a result of the whole-body lean phenotype, despite *Adggr6* lower expression being observed only in pWAT. The *Adgrg6*^*ARS-/-*^ male mice did not show significant difference in GTT and ITT under chow diet (**fig. S5D**). We next placed *Adgrg6*^*ARS-/-*^ male mice on HFD for thirteen weeks (**Fig. 4G**). *Adgrg6*^*ARS-/-*^ mice had significantly lower body weight than littermate control mice (**Fig. 4H**). In addition, these mice had smaller iWAT and pWAT mass (**Fig. 4I**). *Adgrg6*^*ARS-/-*^ mice also showed improved GTT and ITT compared to control males (**Fig. 4J**), indicating that *Adgrg6* enhancer knockout can protect mice against diet-induced obesity and improve obesity-associated insulin resistance. Combined, our data demonstrate that the *Adgrg6_4* enhancer is VAT-specific, has an important function in adipogenesis and leads to gender-specific effects on adipose tissue distribution.

### CRISPRi-mediated *Adgrg6* knockdown prevents high fat diet-induced obesity

To showcase the therapeutic potential of *Adgrg6* downregulation in males, we utilized CRISPRi, targeting either the *Adgrg6* promoter or *Adgrg6_4* enhancer, to knockdown *Adgrg6* expression *in vivo* (**Fig. 5A**). We first designed five gRNAs targeting either the *Adgrg6* promoter or *Adgrg6_4* enhancer, cloned them into an rAAV-mCherry vector and tested them along with dCas-KRAB in mouse 3T3-L1 preadipocytes. We found two gRNAs targeting the promoter and one gRNA targeting the *Adgrg6_4* enhancer to significantly reduce *Adgrg6* mRNA levels by ∼50-70%, as determined by qRT-PCR (**fig. S6A**). The gRNAs along with dCas9-KRAB were packaged into AAV serotype 9, which is known to effectively express transgenes in mouse adipose depots (*77*). AAV9 co-infection of individual gRNA along with dCas9-KRAB into 3T3-L1 cells found all three gRNAs to significantly reduce *Adgrg6* expression (**fig. S6B**).

**Fig. 5.**
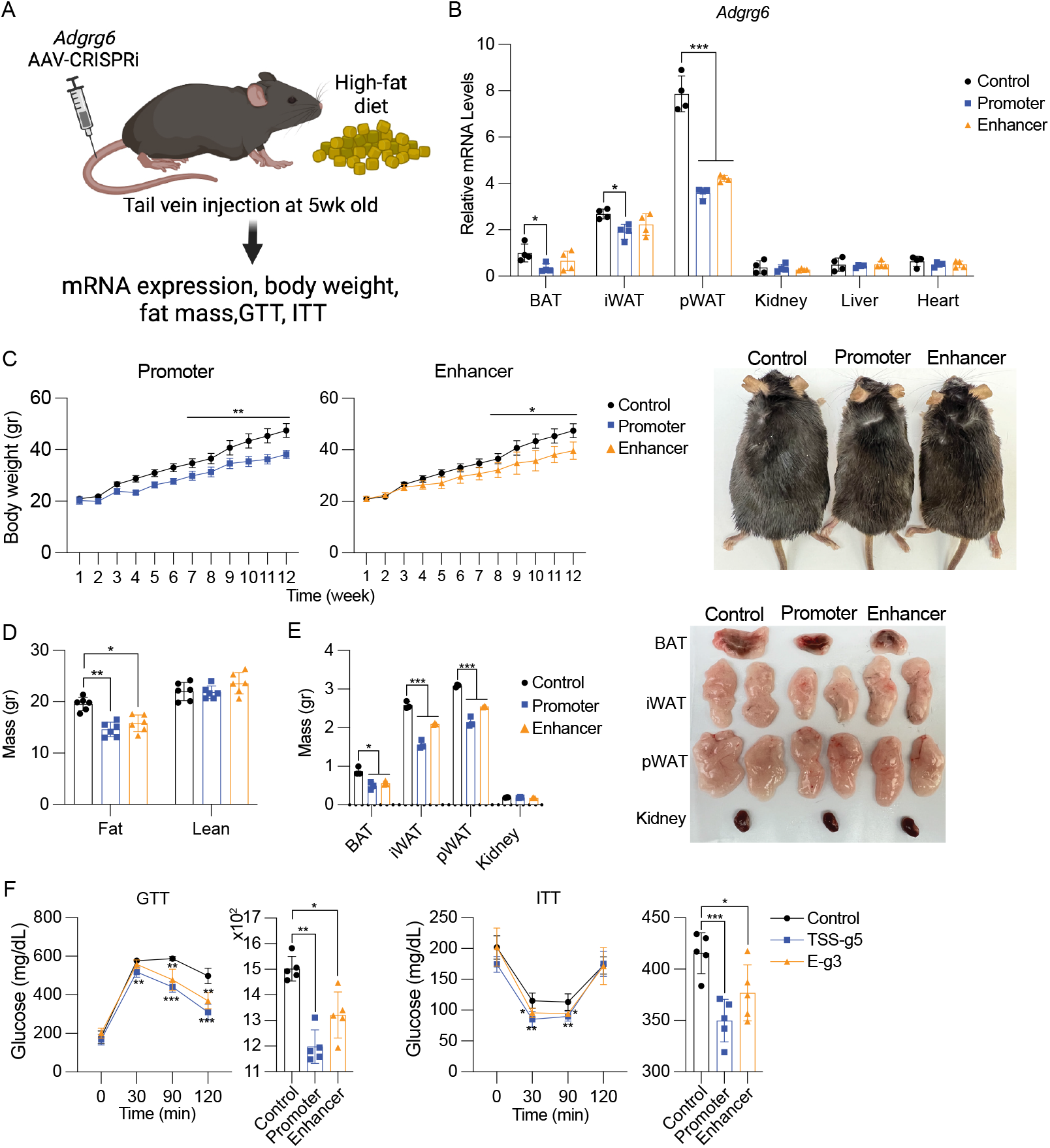
CRISPRi-mediated *Adgrg6* knockdown prevents diet-induced obesity. (**A**) Schematic of AAV-based *Adgrg6* CRISPRi intravenous tail vein injection followed by high-fat diet (HFD) and phenotypic measurements. (**B**) qRT-PCR of *Adgrg6* in brown adipose tissue (BAT), inguinal adipose tissue (iWAT), and perigonadal adipose tissue (pWAT), kidney, liver, and heart of control mice (dCas9-Krab) (n=6), CRISPRi targeting *Adgrg6* promoter (n=6), or enhancer (n=6) mice. Data are represented as mean ± S.D. *≤0.05, ***≤0.001. (**C**) Body weight (left) of control (n=6) and *Adgrg6* promoter (n=6), or enhancer (n=6) AAV-CRISPRi targeted mice on high-fat diet measured for 12 weeks. Data are represented as mean ± S.D. *≤0.05, **≤0.01. Representative mouse images (right) of 20-week-old mice fed on HFD. (**D**) Fat and lean mass of control (n=6), AAV-CRISPRi targeting promoter (n=6), or enhancer (n=6) of *Adgrg6* measured by dual energy X-ray absorptiometry (DEXA). (**E**) Tissue weight (left) of control (n=3), CRISPRi targeting the *Adgrg6* promoter (n=3) or enhancer (n=3). Data are represented as mean ± S.D. *≤0.05, ***≤0.001. Representative tissue images (right) of 20-week-old mice fed on HFD. (**F**) Glucose tolerance test (GTT) with area under the curve (AUC) and insulin tolerance test (ITT) with AUC of control (n=6), CRISPRi targeting promoter (n=6), or enhancer (n=6) of *Adgrg6* of 20-week-old mice fed on HFD. Data are represented as mean ± S.D *≤0.05 **≤0.01, ***≤0.001.

We intravenously injected five weeks old C57BL/6J male mice via tail vein with a guide targeting the *Adgrg6* promoter (g5 which showed the strongest AAV9-based reduction of *Adgrg6* expression) or enhancer (g3) along with another virus encompassing dCas9-KRAB (control). After 2-week of injection, these mice were fed on a HFD for twelve weeks (**Fig. 4A**). We then measured by qRT-PCR, mCherry and *Adgrg6* mRNA levels in adipose tissues, kidney, liver, and heart. mCherry levels were detected in all adipose tissues with the highest level in BAT and pWAT (**Fig. S6D**), consistent with tissue distribution of AAV9 (*77*). *Adgrg6* expression levels were significantly reduced in pWAT, iWAT and BAT in promoter targeted mice and only significantly reduced in pWAT in enhancer targeted mice (the reduction in iWAT and BAT was not significant), fitting with its *Adgrg6* enhancer knockout expression (**Fig. 5B**). These data suggest that dCas9-KRAB CRISPRi, as used in our AAVs, was only able to downregulate gene expression in the tissue where the targeted regulatory element is active, likely due to open chromatin accessibility. CRISPRi *Adgrg6* promoter or enhancer targeted mice showed reduced body weight compared to mice injected with dCas9-KRAB only, as a negative control (**Fig. 5C**). DEXA analyses found these mice also to have lower fat mass compared to dCas9-KRAB only mice (**Fig. 5D**) and smaller BAT, iWAT, and pWAT upon dissection (**Fig. 5E**). The observed lower BAT and iWAT mass in the enhancer targeted CRIPSRi mice could be a result of their overall whole-body lean phenotype. In addition, these mice showed significantly improved GTT and ITT compared to dCas9-KRAB only injected males (**Fig. 5E**). In summary, our data indicate that *Adgrg6* CRISPRi targeting of either the promoter or *Adgrg6_4* enhancer protects mice against HFD obesity and diabetes, suggesting that *Adgrg6* could be a potent therapeutic target to treat these diseases.

## Discussion

Body fat distribution is gender-specific and is a major risk factor for metabolic disease, including obesity, diabetes, and cardiovascular disease. Here, we found that ADGRG6, an adhesive G-coupled receptor, is more highly expressed in visceral fat of males than females. We identified an *ADGRG6* enhancer that overlaps with a SNP that is associated with the visceral fat of females and show that the associated allele leads to reduced enhancer function and lower *ADGRG6* mRNA and protein levels. We further show by either knocking out *ADGRG6*, its enhancer or overexpression that *ADGRG6* promotes adipocyte differentiation by increasing cAMP levels. Adipocyte conditional removal of *Adgrg6* or knocking out its adipocyte enhancer (*Adgrg6_4*) in mice, leads to decreased adiposity and obesity in male mice. These male *Adgrg6*^*ASKO/ASKO*^ or *Adgrg6*^*ARS-/-*^ mice are resistant to HFD-induced obesity with improved glucose tolerance and insulin sensitivity. CRISPRi targeting of the *Adgrg6* promoter or *Adgrg6_4* enhancer protected male mice against HFD-induced obesity and improved insulin response, suggesting *Adgrg6* is a potential therapeutic target to treat obesity and metabolic disease in males.

There are several efforts to map the genetics of favorable adipose phenotypes and elucidate the causal mechanism of gender-specific genetic associations with central obesity and body fat distribution. GWAS provides a powerful tool to identify, in an unbiased manner, loci associated with complex traits and diseases, such as obesity. Large-scale GWAS and whole-exome sequencing efforts have mapped a treasure trove of novel findings for BMI, central obesity phenotypes, adipose distribution, and related metabolic traits.

Interestingly, several studies have found that more than 50% of central obesity loci have significant gender dimorphism (*37, 38, 78, 79*), with the majority of SNPs residing in noncoding regions of the genome, likely associated with gene regulatory elements that reside in these regions. Here, we characterized one of these loci, *ADGRG6*, but there are many more associated loci to identify and characterize. It will also be interesting to assess combinations of these SNPs and how they might affect obesity and polygenic risk scores for this phenotype. Interestingly, rs6570507, is also associated with adolescent idiopathic scoliosis (AIS), which is more common in females compared to males (*80-82*). It will be interesting to analyze whether loci influencing a gender specific effect in a certain phenotype could be used to inform regions that are more likely to have these effects also on another phenotype and how this mechanism evolved to influence multiple phenotypes.

Fat depot distribution differences is the major cause of obesity-related diseases (*7-9*). VAT is highly correlated with obesity, insulin resistance and cardiovascular disease while SAT is shown to be metabolically protective against obesity-related metabolic diseases (*4-6*). The differential location of fat has various functional implications on adipokine production, development of insulin sensitivity, inflammation response, mitochondrial function, lipolysis, and free fatty acid release, all of which differ between adipose depots (*10-12, 14, 15, 83*). Our data showed that *ADGRG6* promotes adipocyte differentiation by activating cAMP in preadipocytes. We also found *ADGRG6* to have gender-differential expression in VAT in humans with VAT of males showing higher expression of *ADGRG6* than females. Previous reports have found that numerous genes are differentially expressed in adipose tissue from obese males and females with very few on the sex chromosomes (*35, 36, 84*). However, there is lack of understanding of how these genes are differentially regulated between genders. Here, we found rs6570507 to be highly associated with female trunk fat and reside in an adipocyte enhancer of *ADGRG6*. We demonstrate that this associated SNP alters the binding sites of multiple transcription factors, such as GR and PR that are involved adipogenesis (*64, 85-89*) and disrupts the binding affinity of these proteins, leading to lower expression of *ADGRG6*. Our work showcases how genetic variation can allow to identify an enhancer that can lead to gender-differential gene expression and phenotype, gender-specific fat distribution.

Ablation of *Adgrg6* in adipose tissue using *Pdgfra*- or *Adipoq*-driven Cre (*Adgrg6*^*ASKO*^ and *Adgrg6*^*Adipoq-Cre*^) or its enhancer (*Adgrg6*^*ARS-/-*^) in mice all led to reduced BW and VAT mass in male mice but not in female mice. Male mice from all three deletions develop less VAT and exhibit similar metabolic phenotypes to female mice. They also demonstrated lower adiposity with improved glucose tolerance and insulin sensitivity compared to WT male mice. Moreover, these male mice also became resistant to high-fat diet-induced obesity. Similar to humans, *Adgrg6* expression was higher in VAT than SAT in male mice compared to female mice. Thus, the gender-specific body weight and adiposity phenotypes we observed in *Adgrg6*^*ASKO/ASKO*^ or *Adgrg6*^*Adipoq-Cre*^ might be due to the fact that *Adgrg6* basal expression level is low in VAT of females compared to males. Deleting this gene or the *Adgrg6_4* enhancer, that is conserved between mouse and humans, did not have a significant impact on VAT development in females. Interestingly, ablation of the *Adgrg_4* enhancer reduced *Adgrg6* mRNA levels only in pWAT (**Fig. 4B**), indicating visceral fat specificity for this enhancer. This was further corroborated by its CRISPRi knockdown which significantly lowered *Adgrg6* expression only in pWAT (**Fig. 5B**). To date, Wilms Tumor 1 (WT1) is the only reported factor to be highly expressed in visceral fat (*90*) and its promoter is used to drive *Cre* to conditionally ablate genes in visceral fat (*91*). The *Adgrg_4* enhancer could also be utilized to drive tissue specific expression of a gene of interest in pWAT of male mice. While further work needs to be done, for example in non-human primates, to suggest this gene and regulatory element have a similar effect in humans, our results provide strong support for its role in adipogenesis and gender specific fat distribution.

*Cis* regulation therapy (CRT), the use of nuclease deficient gene editing systems coupled to transcription modulating proteins has shown great promise to treat genetic disease (*92*). For example, our lab has demonstrated that by upregulating via CRISPRa either *Sim1* or *Mc4r* in heterozygous mice we can rescue their obese phenotype (*93*). Gender-specific fat distribution contributes to differential metabolic outcomes and co-morbidities. Utilizing CRISPRi, we show that downregulation of *Adgrg6* via CRISPRi by targeting either its promoter or enhancer could provide a viable therapeutic option for male obesity and its associated co-morbidities. Our data clearly shows that our gRNAs specifically target *Adgrg6* in adipose tissues and can lower body weight, adiposity, and improved insulin sensitivity. Interestingly, we also observed tissue-specific downregulation depending on the targeted regulatory element. For the promoter targeted mice, we observed downregulation of *Adgrg6* in all examined adipose tissues, while the enhancer targeted mice only showed significant downregulation in pWAT. This is in line with our previous work showing targeted regulatory element tissue-specific upregulation with CRISPRa (*93*), likely due to dCas9 binding being more amendable in low nucleosome occupancy regions, as previously shown for various CRISPR screens (*94, 95*). This regulatory element guided tissue-specificity provides an added advantage for CRT over standard gene replacement therapy. Further development of CRT approaches that specifically target and downregulate *ADGRG6* could provide a novel strategy to combat obesity and its co-morbidities in males. In addition, fat transplantation could also be considered for this treatment. As it is commonly used in many surgical procedures, such as aesthetic and reconstructive surgery, it could readily be used for therapeutic treatments (*96*). In humans, there is a growing interest to carry out human adipose tissue grafting by using adipose stem cells/progenitors due to their resistance to trauma and long-term survival following transplantation (*97-99*). Modulating *ADRG6* in VAT and transplanting it in individuals might be novel way to treat obesity and associated metabolic diseases.

## Materials and Methods

### UK Biobank Cohort and Sex-Specific Analyses at ADGRG6

The UK Biobank (UKB) is a large prospective study following the health of approximately 500,000 participants from five ethnic groups (European, East Asian, South Asian, African British, and mixed ancestries) resident in the UK aged between 40 and 69 years-old at the baseline recruitment visit (*51, 100*). Demographic information and medical history were ascertained through touch-screen questionnaires. Participants also underwent a wide range of physical and cognitive assessments, including blood sampling. In the UKB, the trunk fat mass phenotype data (Data-Field 23128 on the web-site: https://biobank.ndph.ox.ac.uk/showcase/field.cgi?id=23128; description: Trunk fat mass in kg; category: Body composition by impedance ; version March 2022) were collected on 492,768 participants. Phenotyping, genotyping, and imputation were carried out by members of the UK Biobank team. Imputation to the Haplotype Reference Consortium (HRC) has been described (www.ukbiobank.ac.uk), and imputation at a few non-HRC sites (for replication) was done pre-phasing with Eagle (*101*) and imputing with Minimac3 (*102*) with the 1000 Genomes Project Phase I. We first ran a linear regression of trunk fat mass and SNP rs9403383 at *ADGRG6* using PLINK (*103*) v1.9 (www.cog-genomics.org/plink/1.9/) adjusting for age, sex, and ancestry principal components (*104*). We then assessed the association between trunk fat mass and SNP rs9403383 by sex (245,000 women and 205,628 men, respectively, in self-reported whites with global ancestry PC1≤70 and PC2≥80, as described(*104*)). The analyses presented in this paper were carried out under UK Biobank Resource project #14105.

### Human adipocytes

For adipocyte differentiation, human preadipocytes (kind gift from Dr. Hei Sool Sul, UC Berkeley) were cultured to 100% confluency in DMEM, supplemented with 10% FBS and fresh media were replaced. After 48hr, cells were subjected to adipocyte differentiation by adding MesenCult Adipogenenic Differentiation Kit (Stemcell, 05412). Media was replaced every 2 days during differentiation. To generate *ADGRG6* KO, *ADGRG6* EKO, and *ADGRG6* AKI cells, human preadipocytes in a 6-well plate were transfected with 6.25ug Cas9 protein (Fisher Scientific, A36498) and 800ng sgRNAs (IDT), 1.5ug ssDNA donor (IDT) (for *ADGRG6* AKI) and 0.5ug GFP plasmid (Addgene, 13031) using LipoMag transfection reagent (OZ Biosciences, LM80500) following the manufacturer’s protocol. After 48 hours, GFP+ cells were isolated into 96 well-plates into single clones using FACS (BD FACSAria Fusion). These colonies were then genotyped to collect properly edited clones. For *ADGRG6* overexpression, in 12-well plates, sub-confluent human preadipocytes were transfected with 500ng *ADGRG6* expression plasmid Myc-DDK-tagged human G protein-coupled receptor 126 (Origene, RC212889) using LipoMag transfection reagent and cells were then subjected to the adipocyte differentiation protocol described above.

### Luciferase Assay

*ADGRG6_3* and *ADGRG_4* sequences were PCR amplified from human genomic DNA and cloned into the pGL4.23 plasmid (Promega, E8411). Human preadipocytes, in a 12-well plate, were transfected with 0.5ug luciferase constructs and 50ng pGL4.73 [hRluc/SV40] plasmid (Promega, E6911) containing Renilla luciferase, to correct for transfection efficiency, using LipoMag transfection reagent (OZ Biosciences) following the manufacturer’s protocol. Empty pGL4.23 was used as a negative control and pGL4.13 (Promega, E668A) with an SV40 early enhancer as a positive control. Forty-eight hours after transfection, cells were lysed, and luciferase activity was measured using Dual-Luciferase Reporter Assay System (Promega, E1910). Six technical replicates were performed for each condition.

### RNA Isolation and qRT-PCR

Total RNA from sorted or cultured cells was extracted using Trizol reagent (ThermoFisher, 15596026). Reverse transcription was performed with 1μg of total RNA using qScript cDNA Synthesis Kit (Quantabio, 95047) following the manufacturer’s protocol. qRT-PCR was performed on QuantStudio 6 Real Time PCR system (ThermoFisher) using Sso Fast (Biorad, 1705205). Statistical analysis was performed using ddct method with GAPDH primers as control (see primer sequences in **Table S1**). Gene expression results were generated using mean values for over n=4-6 biological replicates.

### Separation of SVF and Adipocyte Fractions

SVF fractionation was carried out as previously described (*103*). Subcutaneous and visceral mouse adipose tissue depots were dissected from 16 week-old C57BL/6J mice into ice-cold PBS. The tissue was finely minced using scissors and digested with Collagenase type II (Sigma-Aldrich, C2-22)in 3% BSA-HBSS buffer at 37°C for 45 minutes with gentle shaking. The cell suspension was then centrifuged. The floating fraction containing adipocytes were collected and the pellet was resuspended and passed through 100μm cell strainer and spun at 500g for 5 minutes. The cell pellet was resuspended in HBSS buffer and passed through 70μm and 40μm cell strainers to ensure a single cell preparation. The cells were spun at 500g for 5 minutes to pellet, and the red blood cells in the pellet were then lysed by incubating the pelleted cells in red blood cell lysis buffer on ice for 5 min with shaking. The remaining cells were washed twice with HBSS and spun at 500g for 5 min to pellet. SVF cells were incubated with indicated antibodies for 20 minutes in the dark, washed, spun at 300g for 5 minutes, resuspended in PBS and passed through a 40μm filter prior to FACS analysis. FACS was performed on ARIA Fusion Cell Sorter. Cells that were not incubated with antibody were used as a control to determine background fluorescence levels. Cells were initially chosen based on forward and side scatter (FFS, and CCS). Single cells were then separated into Lin+ cells (CD45+, CD31+, Ter119) and Lin-cells. In the Lin-fraction, messenchymal stem cells (MSC) were then selected by CD109, adipose progenitors (APC) were sorted by CD34 and Pdfrga (indicated anitbodies in **Table S2**). Cells were collected in PBS for RNA isolation or in DMEM+20% FBS for cell culture.

### Oil red O staining

Differentiated adipocytes were fixed with 1% paraformaldehyde (PFA) for 20 minutes. Cells were then washed 3x with water and incubated in 60% isopropanol. Oil red O staining solution (3mg/ml) was added to the cells and followed by staining for 15 minutes. Cells were then washed with water and images were obtained with a Leica DFC700T. Lipid droplet size and number were analyzed by Fiji software (*59*). At minimum of 100 cells were counted for each condition.

### ChIP qPCR

ChIP was performed using ChIP-IT Express Chromatin Immunoprecipitation kits (Active Motif, 53009). Briefly, cells cultured on 15cm plate were harvested and fixed with 1% PFA for 10 minutes. The reaction was stopped by incubating with 125 mM glycine for 10 minutes. Cells or tissues were rinsed with ice-cold phosphate-buffered saline (PBS) for three times and lysed in IP lysis buffer. Nuclei were collected by centrifugation at 600 × g for 5 minutes at 4°C. Nuclei were released by douncing on ice and collected by centrifugation. Nuclei were then lysed in nuclei lysis buffer and sonicated three times by 20 second bursts, each followed by 1 minute cooling on ice using Covaris m220 ultrasonicator. Chromatin samples were diluted 1:10 with the dilution buffer containing 16.7 mM Tris pH 8.1, 0.01% SDS 1.1% Triton X-100 1.2 mM EDTA, 1.67 mM NaCl, and proteinase inhibitor cocktail. Antibodies (2ug) of Histone H3 (Abcam, ab4729), glucocorticoid receptor (Abcam, ab3671), progesterone receptor (Abcam, ab2765), and HoxA3 (Sigma-Aldrich, H3791) or normal mouse IgG (Diagenode, C15400001) and protein A/G magnetic beads (Thermo-Fisher, 88802) were used. After the antibody-bead-chromatin mixtures were incubated at 4C overnight, the beads were washed and cross-linked reversed. DNA fragments were purified, and samples were analyzed by qPCR for enrichment in target area of *Adgrg6* enhancer using primers targeting 150bp around the associated SNP (**table S1**). Four replicates were used for each antibody. The fold enrichment values were normalized to input.

### Mouse lines

All animal studies were carried out in accordance with University of California San Francisco Institutional Animal Care and Use Committee. Mice were housed in a 12:12 light-dark cycle, and chow diet (Envigo, 2018S) and water were provided *ad libitum*. Mice were fed with either chow diet or high fat diet with 40% fat (Research Diet, D12492i). *GPR126*^*fl/fl*^ mice (*64*) were provided by Dr. Ryan Gray. *Adgrg6*^*ASKO/ASKO*^ constitutive adipose tissue knockout mouse were generated by cross-mating *Pdgfra*-driven Cre mouse (Jackson Laboratory, 012148) and *GPR126*^*fl/fl*^ mouse. *Adgrg6*^*Adipoq-Cre*^ mice were created by crossing the *Adipoq*-driven *Cre* mouse (Jackson Laboratory, 010803) with the *GPR126*^*fl/fl*^ mouse. The *Adgrg6*^*ARS-/-*^ mouse was generated as described in (*104*). Briefly, the 4 kilobase *Adgrg6* adipose regulatory sequence was converted to mouse sequence using the UCSC Genome Browser LiftOver tool (*73*). Two gRNAs (IDT) (30uM), designed to target the 5′ and 3′ ends of this region (**table S1**), were mixed with Cas9 (1mg/ml) protein and were injected into the oviduct lumen of female mice that were mated the night before. Subsequent offspring were genotyped via PCR-Sanger sequencing of the breakpoint and Southern blot. PCR-Sanger sequencing was preformed using standard techniques using three primers (**table S1**). For Southern blot analyses, genomic DNA was treated with *Avr*II (New England Biolabs, catalog no. R0193) and fractionated by agarose gel electrophoreses. Following capillary transfer onto nylon membranes, blots were hybridized with Digoxigenin (DIG)-labeled DNA probes (corresponding to chr2:147,202,083-147,202,444; mm9) ampli?ed by the PCR DIG Probe Synthesis Kit (Sigma-Aldrich, 11636090910). The hybridized probe was immunodetected with antidigoxigenin Fab fragments conjugated to alkaline phosphatase (Sigma-Aldrich, 11093274910) and visualized with a CDP star (Sigma-Aldrich, 11685627001) according to the manufacturer’s protocol. Chemiluminescence was detected using the FluorChem E (ProteinSimple, 92-14860-00).

### Body composition and food intake analyses

Body composition was measured using dual energy x-ray absorptiometry (DEXA) by PIXImus Mouse Desnitometer (GE Medical Systems). For DEXA, mice were anesthetized with 2% isoflurane and measured for bone mineral density and tissue composition (fat mass and lean mass). Food intake was measured by using the Columbus Instruments Comprehensive Lab Animal Monitoring System (CLAMS) (Columbus Instruments). Mice were housed individually and acclimatized on powdered picodiet (PicoLab 5058) for 3 to 4 days, and food intake measurements were done over 4 to 5 days.

### Glucose tolerance test (GTT)

Mice were fasted overnight. Blood glucose of each mouse was measured to get the value for the 0 time point. Blood glucose level was determined by Contour Next Blood Glucose meter (Contour, 7277) in tail vein blood. Tail snipping was used to get blood. Glucose (1g/kg of BW) (Sigma-Aldrich, G8270) was then administered intraperitoneally. Blood glucose was measured at 30, 60, and 120 minutes after injection.

### Insulin tolerance test (ITT)

Mice were fasted for 4 hours. Blood glucose was measured for each mouse to get the value for the 0 time point. Blood glucose levels were determined by Contour Next Blood Glucose meter (Contour, 7277) in tail vein blood. Recombinant human insulin (0.75 IU insulin/kg BW) (Sigma-Aldrich, I9278) was then administered intraperitoneally. Blood glucose was measured at 30, 60, and 120 minutes after injection.

### CRISPRa AAV *in vitro* optimization

Five gRNAs targeting the promoter of human *ADGRG6* or mouse *Adgrg6* were designed using the Broad Institute CRISPick Tool (*105*) (**table S1**). These guides were individually cloned into pAAV-U6-sasgRNA-CMV-mCherry-WPREpA (*92*) at the *BstXI* and *XhoI* restriction enzyme sites using the In-Fusion (Takara Bio, 638910) cloning methods as described in (*92*). Mouse gRNA constructs and pCMV-sadCas9-KRAB (kind gift from Dr. Alejandro Lomniczi OHSU) were co-transfected into human preadipocytes X-tremeGene (Sigma, 6366236001) while human gRNA plasmids and pCMV-sadCas9-KRAB were co-transfected into mouse 3T3-L1 preadipocytes. Both cell lines were maintained in DMEM supplemented with 10% FBS. After 48 hours, cells were lysed with Trizol and RNA was collected and cDNA and qRT-PCR were performed, as mentioned for RNA Isolation and qRT-PCR, and differential expression was determined using ddct method with GAPDH primers as control. The two gRNAs that reduced human *ADGRG6* or mouse *Adgrg6* expression the most were packaged into rAAV-9 serotype virons. rAAV-9 serotype virons were produced by transfecting AAVpro 293T cell (Takara, 6322723) with pCMV-sadCas9-KRAB (Addgene, 115790) or pAAV-U6-sasgRNA-CMV-mCherry-WPREpA (*92*) along with packaging vectors, including PAAV2/9n (Addgene, 112865) and pHelper vectors using TransIT293 reagent (Mirus, 2700). After 72 hours, AAV particles were collected and purified using AAVpro Cell & Sup. Purification Kit Maxi (Takara, 6676) and quantified by the AAVpro Titration Kit (Takara, 6233). gRNA AAV viruses (1×10^3^ MOI) and dCAS9-KRAB AAV (1×10^3^ MOI) were used to infect human or mouse preadipocytes. After five days, RNA was collected, and cDNA was made and qRT-PCR was done as previously described. Differential expression was determined using the ddct method with GAPDH primers as control (primer sequences in **table S1**).

### CRISPRi mouse injections

C57BL/6J mice at 4wk old were kept under live anesthetic isoflurane at 1.0%-2.0%. A single dose of AAV9-dCas9-Krab and AAV9-gRNA (1×10^11^vg/mouse) at 1:1 ratio were injected intravenously via tail-vein. Body weight was recorded weekly. Five-week post-injection, mice were subjected to DEXA, GTT, and ITT.

## Acknowledgments

We would like to thank Dr. Hei Sook Sul for kindly providing the human preadipocyte cell line and Dr. Alejandro Lomniczi for kindly providing providing the pCVM-sadCas9-KRAB vector.

## Funding

This work was funded in part by:

National Institute of Diabetes and Digestive and Kidney Disease (NIDDK) R01DK116738 (H.C. and N.A) and 1R01DK124769 (N.A)

National Human Genome Research Institute (NHGRI) grant number R01HG012396 (N.A)

California Institute for Regenerative Medicine (CIRM) postdoctoral fellowship.

## Author contributions

Conceptualization: HPN, NA

Methodology: HPN, AU, RS, KA, CB, HC, RG, NA

Formal analysis: HPN; Investigation-HN, NA

Writing, Review & Editing: HPN, NA

Visualization: HPN, NA

Supervision: NA

Funding acquisition: NA

## Data and materials availability

All data and analysis code are available upon request. Requests for materials can be directed to NA (CRT-based approaches).

## Supplementary Materials

Materials and Methods

Figures S1 to S6

Tables S1 to S2

