## Supplementary material for "ADGRG6 promotes adipogenesis and is involved in sex-specific fat distribution": Supplmentary materials

A

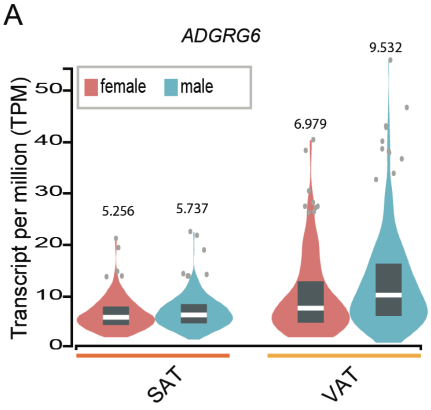


B

| UK BioBank association | | | |
|---|---|---|---|
| Trait | Sex | p-value | Source DB |
| **Trunk fat mass** | f & m | $2.57 \times 10^{-11}$ | UK Biobank |
| **Trunk fat mass** | F | $2.61 \times 10^{-11}$ | UK Biobank |

C

| Trait | Probe | Tissue | Statistic (p-value) | Source |
|---|---|---|---|---|
| ***ADGRG6*** | ENSG00000112414 | subcutaneous adipocytes | $5.2 \times 10^{-13}$ | GTEX Portal V8 |
| ***ADGRG6*** | ENSG00000112414 | unexposed skin | $3.05 \times 10^{-11}$ | GTEX Portal V8 |
| ***ADGRG6*** | ENSG00000112414 | sun exposed skin | $6.13 \times 10^{-8}$ | GTEX Portal V8 |
| ***ADGRG6*** | ENSG00000112414 | visceral adipocytes | $3.26 \times 10^{-5}$ | GTEX Portal V8 |
| ***ADGRG6*** | ENSG00000112414 | skeletal Muscle | $3.68 \times 10^{-5}$ | GTEX Portal V8 |
| ***ADGRG6*** | ENSG00000112414 | muscularis mucosae | $8.14 \times 10^{-5}$ | GTEX Portal V8 |

**Fig. S1. *ADGRG6* is highly expressed in male visceral fat.** (**A**) *ADGRG6* expression in human subcutaneous adipose tissue (SAT) and visceral adipose tissue (VAT) of men and women obtained by GTEX portal. (**B**) UK Biobank association data for rs9403383 in trunk fat mass. (**C**) *cis*-eQTL results for *ADGRG6* locus in various tissues.

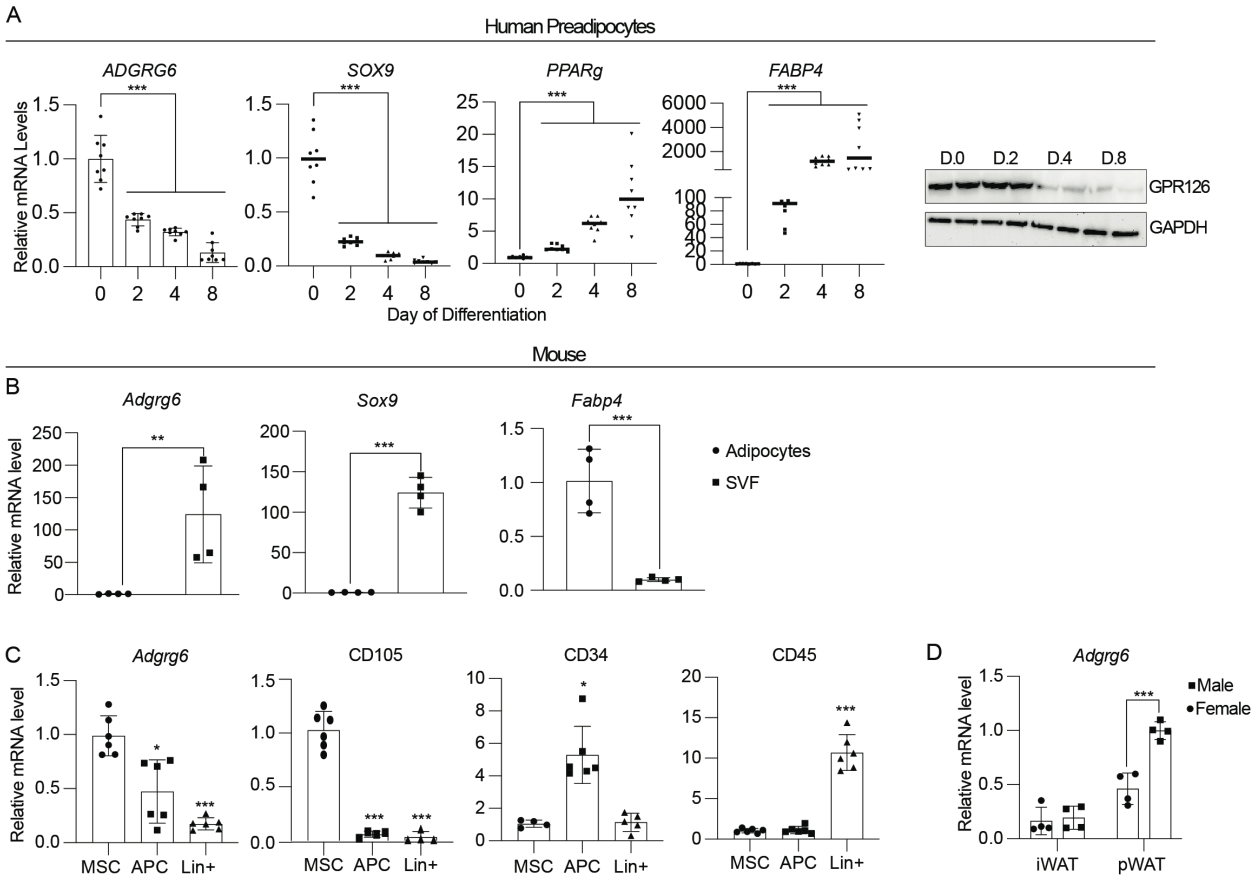


**Fig. S2.** ***ADGRG6* is highly expressed in preadipocytes and decreased upon adipocyte differentiation.** (**A**) qRT-PCR of *ADGRG6*, *SOX9*, *PPARg*, and *FABP4* during adipocyte differentiation of human preadipocytes (left). Data are represented as mean ± S.D. ***≤0.001. Immunoblotting of ADGRG6 and GAPDH during human adipocyte differentiation (right). (**B**) qRT-PCR of *Adgrg6*, *Sox9*, and *Fabp4* in adipocytes and stromal vascular fraction (SVF) in mouse perigonadal white adipose tissue (pWAT). Data are represented as mean ± S.D. **≤0.01, ***≤0.001. (**C**) qRT-PCR of *Adgr6*, *CD109*, *CD34*, and *CD45* in mesenchymal stem cells (MSC), adipose progenitor cells (APC), and linage positive cells (Lin+), which were FACS-sorted from mouse pWAT. Data are represented as mean ± S.D. *≤0.05, ***≤0.001. (**D**) qRT-PCR of *Adgr6* in mouse inguinal white adipose tissue (iWAT), and pWAT of male and female C57BL/6J mice. Data are represented as mean ± S.D. ***≤0.001.

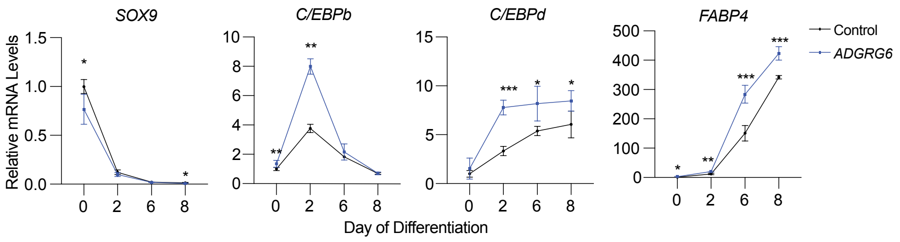


**Fig. S3. *ADGRG6* overexpression increases expression of adipogenic genes.** qRT-PCR of *SOX9*, *C/EBPb*, *C/EBPd*, and *FABP4* in human adipocytes overexpressing *ADGRG6* during adipocyte differentiation. Data are represented as mean ± S.D. *≤0.05, **≤0.01, ***≤0.001.

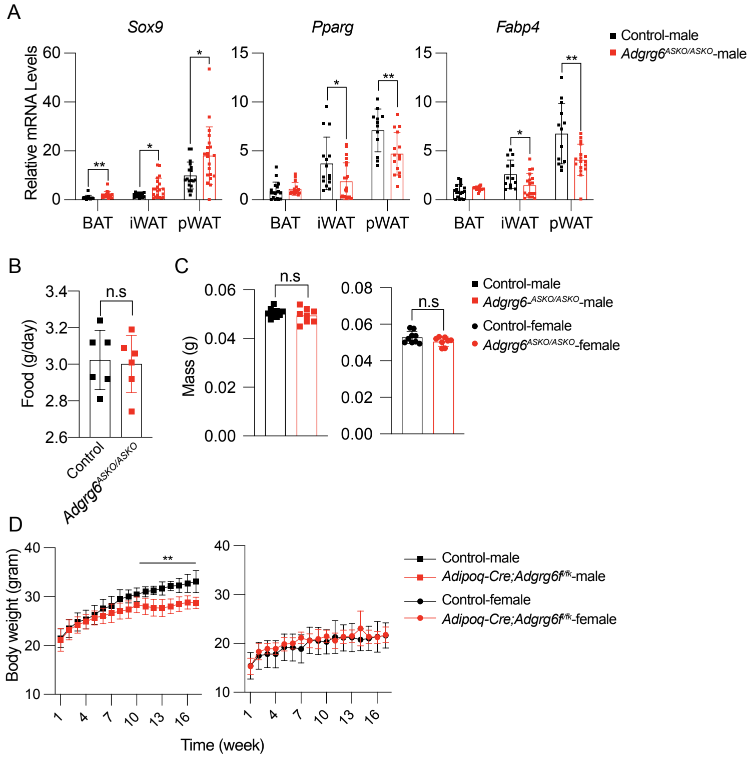


**Fig. S4. *Adgrg6* adipose-specific knockout results in decreased expression of adipogenic genes in adipose tissues.** (**A**) Food intake of male control (N=6) and *Adgrg6*$^{ASKO/ASKO}$ (N=6) mice measured by CLAMS (Comprehensive Lab Animal Monitoring System). (**B**) Bone mineral density of control *Adgrg6*$^{fl/fl}$ (male=9 mice, female=9) and *Adgrg6*$^{ASKO/ASKO}$ (male=8, female=8) by dual energy X-ray absorptiometry (DEXA). (**C**) qRT-PCR of *Sox9*, *Pparg*, and *Fabp4* male control and *Adgrg6*$^{ASKO/ASKO}$ mice. Data are represented as mean ± S.D. *≤0.05, **≤0.01. (**D**) Body weight of *Adipoq*$^{Cre}$ (male=6, female=4) and *Adgrg6*$^{Adipoq\text{-}Cre}$ (male=6, female=4) mice. Data are represented as mean ± S.D. **≤0.01.

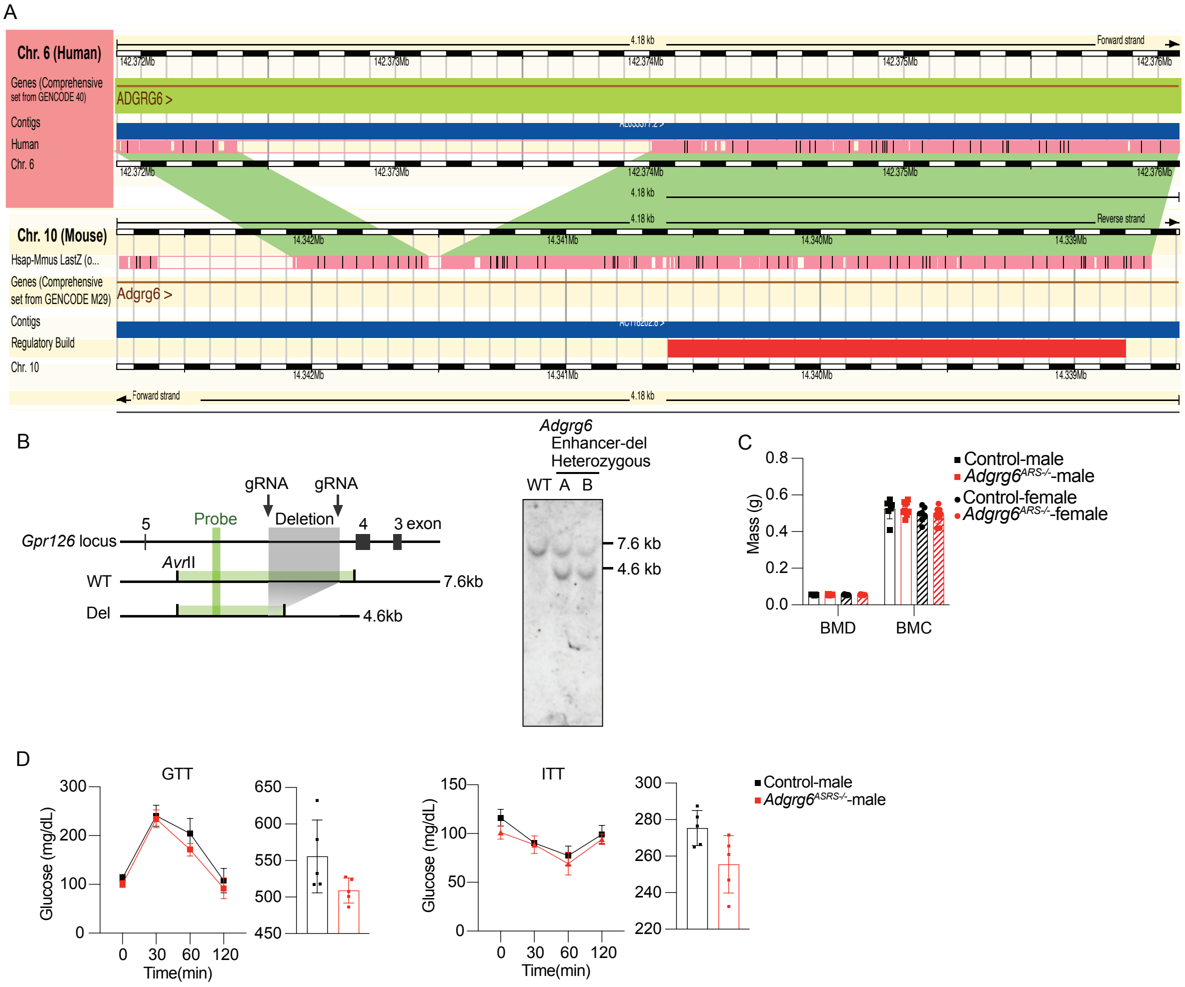


**Fig. S5.** ***Adgrg6* enhancer knockout generation.** (**A**) Genomic alignment of *Adgrg6_4* enhancer in both humans and mice using ENSEMBL. (**B**) Schematic of the genomic location of the *Adgrg_4* enhancer and *Avr*II restriction enzyme sites (vertical lines) in wild type (WT) and deletion (Del) loci. The deletion and probe location are shown as a gray and green rectangle respectively. Expected band sizes in WT and KO are written to the right of the map. Southern blot analyses of wild type (WT) and heterozygous mice (line A and B) with the estimated band size written to the right of the blot. (**C**) Bone mineral density (BMD) and bone mineral content (BMC) of control and *Adgrg6*$^{ARS-/-}$ male and female mice. (**D**) Glucose tolerance test (GTT) and insulin tolerance test (ITT) of male control and *Adgrg6*$^{\mathrm{ARS-/-}}$ mice with area under the curve (AUC) to the right of each graph.

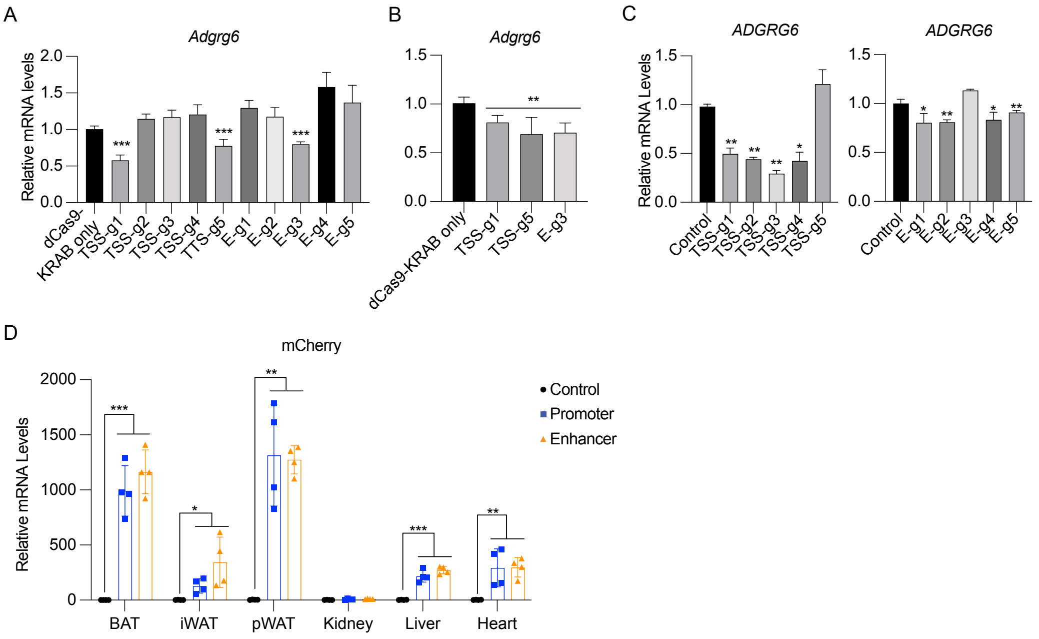


**Fig. S6. CRISPRa AAV *in vitro* optimization.** (**A**) qRT-PCR of *Adgrg6* in mouse 3T3-L1 preadipocytes transfected with dCas9-KRAB and various gRNAs targeting the promoter or the enhancer of *Adgrg6*. Data are represented as mean ± S.D. ***≤0.001. (**B**) qRT-PCR of *Adgrg6* in mouse 3T3-L1 preadipocytes infected with AAV9-dCas9-KRAB and two gRNAs targeting the promoter or one gRNA targeting the enhancer of *Adgrg6*. Data are represented as mean ± S.D. **≤0.01. (**C**) qRT-PCR of *ADGRG6* in human preadipocytes transfected with dCas9-KRAB and various gRNAs targeting the promoter or the enhancer of *ADGRG6*. Data are represented as mean ± S.D. *≤0.05, **≤0.01. (**D**) qRT-PCR of mCherry in adipose tissues, kidney, liver, and heart of control, CRISPRi targeting the promoter or enhancer of *Adgrg6* mice. Data are represented as mean ± S.D. *≤0.05, **≤0.01, ***≤0.001.

| Gene Symbol | Primers |
| --- | --- |
| *Gapdh* | TGC ACC ACC AAC TGC TTA G |
| | GGA TGC AGG GAT GAT GTT C |
| *Adgrg6* | TCC TGT CCA TCT CTG GCT CA |
| | CAC AAG ACA GAG CTG CTC CA |
| *Sox9* | CAC ACG TCA AGC GAC CCA TGA A |
| | TCT TCT CGC TCT CGT TCA GCA G |
| *Pparg* | GTA CTG TCG GTT TCA GAA GTG CC |
| | ATC TCC GCC AAC AGC TTC TCC T |
| *C/ebpb* | CAA CCT GGA GAC GCA GCA CAA G |
| | GCT TGA ACA AGT TCC GCA GGG T |
| *C/ebpd* | AAA GTG CAG GCT TGT GGA CT |
| | TTA CTC CAC TGC CCA CCT GT |
| *Fabp4* | GCT TGT CAC CAT CTC GTT TTC TC |
| | TGA AAT CAC CGC AGA CGA CAG G |
| *GAPDH* | GTC TCC TCT GAC TTC AAC AGC G |
| | ACC ACC CTG TTG CTG TAG CCA A |
| *SOX9* | AGG AAG CTC GCG GAC CAG TAC |
| | GGT GGT CCT TCT TGT GCT GCA C |
| *PPARg* | AGC CTG CGA AAG CCT TTT GGT G |
| | GGC TTC ACA TTC AGC AAA CCT GG |
| *C/EBPb* | AGA AGA CCG TGG ACA AGC ACA G |
| | CTC CAG GAC CTT GTG CTG CGT |
| *C/EBPd* | TCC GGC AGT TCT TCA AGC AGC T |
| | GAG GTA TGG GTC GTT GCT GAG T |
| *FABP4* | ACG AGA GGA TGA TAA ACT GGT GG |
| | GCG AAC TTC AGT CCA GGT CAA C |
| CD34 | GTC AAG TTG TGG TGG GAA GA |
| | AGA GGC GAG AGA GGA GAA AT |
| CD45 | ACA TCA TCG CCAGCATCTATC |
| | CTT GCC TCC ATC CAC TTC ATT A |
| CD90 | CCT TAC CCT AGC CAA CTT CAC C |
| | TTA TGC CGC CAC ACT TGA CCA G |
| *ADGRG6* enhancer genotyping | AAT TTG AGC CCC CTC AAA GAG T |
| | GCT CCT TTA GCA TCA GAA AAC CT |
| | AGG GTA GCA TTT TGG TCA GGA |
| *ADGRG6* enhancer ChIP | TTC AGG GGT CTC TTT TCA CCA |
| | AAC CCG TAA CAT GGG ATC TAT GG |
| *ADGRG6_3* | TTC AAA TAC TGT GCT GTC TCT GG |
| | GCT CAT ATT TAG CAG ACATTGTCCA |
| *ADGRG6_4* | TCC TGA GGG AGT GGT ACA GT |
| | CAG GTG AAC TAA CAT TGT GAA CT |
| *Adgrg6* TSS g1* | GGC CCC AGC TCT AGG CGT TCA GAG GA |
| *Adgrg6* TSS g2 | GTG GCC CCA GCT CTA GGC GTT CAG AG |
| *Adgrg6* TSS g3 | AGG TGC GGC TCG GCG CGC CCC CCG GG |
| *Adgrg6* TSS g4 | CAG CTC TAG GCG TTC AGA GGA CAG AG |
| *Adgrg6* TSS g5* | CGC CTA GAG CTG GGG CCA CCC GGG GG |
| *Adgrg6* enhancer g1 | AGT TAA TGA GAA TCG ACT GCA TTG AG |
| *Adgrg6* enhancer g2 | ACA GGC ACA GTA GTA AGA CAT CTG GG |

| | |
|---|---|
| *Adgrg6* enhancer g3* | ATT GGA GAA AAT CCT CTA CAC CAG GG |
| *Adgrg6* enhancer g4 | AAC AACA GGG CCA AAT TGT CTG AG AG |
| *Adgrg6* enhancer g5 | GGG ACT AGA GAT TAT CAA CCA AAG GG |
| *ADGRG6* TSS g1* | AGC TGA GGA AGT AGG GTG TGC GTG GG |
| *ADGRG6* TSS g2* | GGC GGC AGG TCC CTC CTC GCA GGG AA |
| *ADGRG6* TSS g3* | CCC TCC TCG CAG GGA AGT TGG CAG GG |
| *ADGRG6* TSS g4* | CCT CGC AGG GAA GTT GGC AGG GTG AG |
| *ADGRG6* TSS g5 | CCC TGC CAA CTT CCC TGC GAG GAG GG |
| *ADGRG6* enhancer g1* | CTA CAC CAT AGA TCC CAT GTT ACG GG |
| *ADGRG6* enhancer g2* | GAA TGGTGCACTTCCTGCATTCTGGG |
| *ADGRG6* enhancer g3 | CTG GGT GAC TGT CTT TAA TTC AGG GG |
| *ADGRG6* enhancer g4* | GTC TAT GGTGAA AAG AGA CCC CTG AA |
| *ADGRG6* enhancer g5* | GGA GCT TGG AAA GTT TGG TAT TAG AG |

**Table S1.** Primers for rt-qPCR and subcloning

| Name | Vendor | Catalog number |
|---|---|---|
| Histone H3 | Abcam | ab4729 |
| Glucocorticoid receptor | Sigma-Aldrich | H3791 |
| Progesterone receptor | Abcam | ab2765 |
| Mouse IgG | Diagenode | C15400001 |
| Pparg | Diagenode | C15410367 |
| Gpr126 | Abcam | Ab229583 |
| CD105 | Fisher Scientific | FAB1320G100 |
| CD31 | VWR | 102408 |
| CD90 | Fisher Scientific | BDB561969 |
| CD45 | VWR | 103106-BL |
| Ter 119 | Fisher Scientific | BDB561071 |
| CD34 | Biolegend | 343504 |
| CD140a (PDGFRa) | Fisher Scientific | 11-1401-82 |

**Table S2.** List of antibodies.
